# Using influence measures to test normative use of probability density information derived from a sample

**DOI:** 10.1101/2023.02.05.527165

**Authors:** Keiji Ota, Laurence T Maloney

## Abstract

Bayesian decision theory (BDT) is frequently used to model normative performance in perceptual, motor, and cognitive decision tasks where the outcome of each trial is a reward or penalty that depends on the subject’s actions. The resulting normative models specify how decision makers should encode and use information about uncertainty and value – step by step – in order to maximize their expected reward. When prior, likelihood, and posterior are probabilities, the Bayesian computation requires only simple arithmetic operations: addition, etc. We focus on visual cognitive tasks where Bayesian computations are carried out not on probabilities but on (1) *probability density functions* and (2) these probability density functions are derived from *samples*. We break the BDT model into a serries of computations and test human ability to carry out each of these computations in isolation. We test three necessary properties of normative use of pdf information derived from a sample – *accuracy*, *additivity* and *influence*. Influence measures allows us to assess how much weight *each point* in the sample is assigned in making decisions and allows us to compare normative use (weighting) of samples to actual, point by point. We find that human decision makers violate accuracy and additivity systematically but that the cost of failure in accuracy or additivity would be minor in common decision tasks. However, a comparison of measured influence for each sample point with normative influence measures demonstrates that the individual’s use of sample information is markedly different from the predictions of BDT. We demonstrate that the normative BDT model takes into account the geometric symmetries of the pdf while the human decision maker does not. A heuristic model basing decisions on a single extreme sample point provided a better account for participants’ data than the normative BDT model.

**Author Summary:** Bayesian decision theory (BDT) is used to model human performance in tasks where the decision maker must compensate for uncertainty in order to to gain rewards and avoid losses. BDT prescribes how the decision maker can combine available data, prior knowledge, and value to reach a decision maximizing expected winnings. Do human decision makers actually use BDT in making decisions? Researchers typically compare overall human performance (total winnings) to the predictions of BDT but we cannot conclude that BDT is an adequate model for human performance based on just overall performance. We break BDT down into elementary operations and test human ability to execute such operations. In two of the tests human performance deviated only slightly (but systematically) from the predictions of BDT. In the third test we use a novel method to measure the *influence* of each sample point provided to the human decision maker and compare it to the influence predicted by BDT. When we look at what human decision makers do – in detail – we find that they use sensory information very differently from what the normative BDT observer does. We advance an alternative non-Bayesian model that better predicts human performance.

## Introduction

Bayesian Decision Theory (Wald, 1950; Blackwell & Girshick, 1954; Berger, 1985; Ma, 2019; Maloney & Zhang, 2010) is used to model decision and action selection in a wide variety of experimental tasks (perception: Green & Swets, 1966/1974; Knill & Richards, 1996; visual estimation: Battaglia et al., 2007; Warren et al., 2012; movement planning: Trommershäuser et al., 2003a; 2003b; 2008; Ota et al., 2015; 2016; 2019; motor learning: Körding & Wolpert, 2004; obstacle avoidance: Hudson et al., 2012; hand-eye coordination: Zhang et al, 2012; information sampling: Juni et al., 2016; temporal order judgment: Miyazaki et al., 2006). The BDT model allows us to compare actual human performance against normative performance maximizing expected value. The pattern of deviations between human and normative gives us insight into human cognition, perception and motor planning.

But demonstrating that *overall* human decision making performance (amount of reward earned) approaches that of a normative BDT decision maker does not prove that human observers are mimicking the Bayesian computations in detail. Other decision rules (‘heuristics’) can mimic Bayesian performance arbitrarily closely (Maloney & Mamassian, 2009; Zhang et al., 2013, 2015; Gigerenzer, 2011; Rahnev & Denison, 2018). To fully test the claim that humans are carrying out Bayesian computations, we must also look at performance in detail, trial by trial, as we do here.

An advantage of BDT is that it applies to perceptual, motor, and cognitive tasks where uncertainty is captured by the *probability density function* (pdf) of a continuous random variable. A representative task is shown in Figure 1AB. The decision maker may choose to make a speeded reaching movement to any aim point on a display screen with an irregular, white target region, ***T***. If the decision maker hits within the region he receives a monetary reward, otherwise nothing. The end point ***E* = (*E^x^*, *E^y^*)** where the decision maker touches the screen will differ from the aim point ***A* = (*A^x^*, *A^y^*)** because of the motor uncertainty inherent in speeded movements.

**Figure 1.**
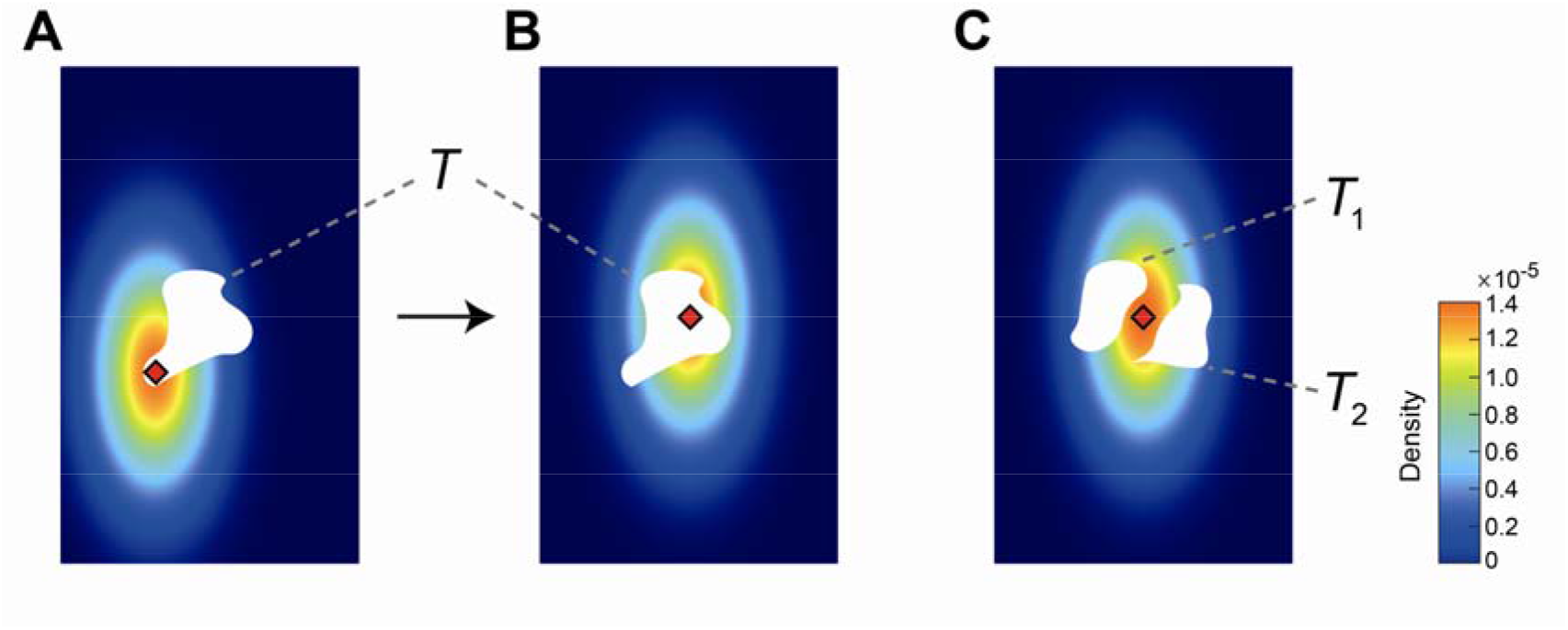
Applications of Bayesian decision theory. **A**. The decision maker is rewarded for making a speeded reaching that terminates in the white target region *T*. He may choose any aim point (red diamond). Because of motor uncertainty his actual end point ***E* = (*E_x_*, *E_y_*)** is distributed as a bivariate Gaussian centered on the aim point, represented here as a heat map. **B**. The decision maker shifts his aim point and the bivariate Gaussian distribution shifts with respect to the aim point. The probability of hitting the target is larger in **B** than in **A**. **C**. The target region is divided into two disjoint regions, ***T*_1_** and ***T*_2_**. Touching either target earns a reward. The decision maker’s aim point is shown in red. Bayesian decision theory allows us to calculate the aim point that maximizes the probability of reward (Trommershäuser et al, 2008).

Where should the decision maker aim? Because the movement is speeded, the outcome of the movement is distributed as a continuous random variable with *probability density function* (*pdf*) ***ϕ*(*E^x^*, *E^y^*|*A*)**. This *population pdf* in reaching tasks is typically close to bivariate Gaussian (Trommershäuser et al., 2003a; 2003b; 2008). In Figure 1A, a pdf is plotted as a heat map, the aim point as a red diamond, and the target region is in white. The pdf is not itself a probability but it serves to assign a probability that the next reaching movement toward the aim point would end within ***T***:

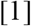

In Figure 1B, we plot the target and pdf for an alternative aim point, also marked in red. Small shifts of the aim point are equivalent to rigidly translating the pdf.^1^

The probability that the decision maker hits the target is plausibly higher in Figure 1B than in Figure 1A. In maximizing the probability of hitting the target (Eq. 1) the decision maker must in effect choose not just between these two aim points but among all possible aim points. The normative BDT decision maker – maximizing expected value – selects the aim point that maximizes Eq. [1] for any choice of target region (Trommershäuser et al., 2003a; 2003b; 2008; for review, see Maloney & Zhang, 2010).

### Accuracy

We refer to the decision maker’s ability to correctly evaluate Eq. [1] for any target as ***accuracy***. The decision maker’s estimates of probability can be accurate for all possible targets only to the extent that he has an accurate estimate of the Gaussian pdf^2^ (Maloney & Mamassian, 2009) or, equivalently the parameters, that determine the Gaussian pdf, the population mean ***μ* = (*μ_x_*, *μ_y_*)** and the population covariance 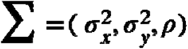. Intuitively, the population mean, ***μ***, specifies the location of the pdf on the plane and the population covariance, ***Σ***, its size and “shape”.

### Additivity

A second task is illustrated in Figure 1C. The target region is now divided into two disjoint regions ***T*_1_** and ***T*_2_** (i.e. ***T*_1_ ∩ *T*_2_ = Ø)**. A reaching movement ending in *either* region earns the same, fixed reward and, as the regions are disjoint, the normative BDT decision maker should seek to maximize

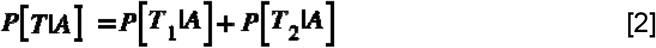

where ***T* = *T*_1_ ∪ *T*_2_**. We refer to the decision maker’s ability to correctly evaluate Eq. [2] as *additivity*. In the first part of this article we compare human estimation with pdfs in simple tasks to that of normative BDT.

#### Decisions based on samples

In many tasks the decision maker does not know the exact population pdf (Figure 2A). The decision maker is instead given a random sample from the population pdf **(*P*_1_, *P*_2_, …, *P_N_*)** as shown in Figure 2B. The sample may be based on prior experience, often provided in an explicit training phase (Maloney & Zhang, 2010). The first question we address is, what information from the sample is retained and used by the decision maker in making any decision? How are the different points in the sample combined to reach a decision? We refer to this information as the **summary** (Figure 2C). The summary is the information about the pdf available to the decision maker and any decision is based solely on this summary (Figure 2D).

**Figure 2.**
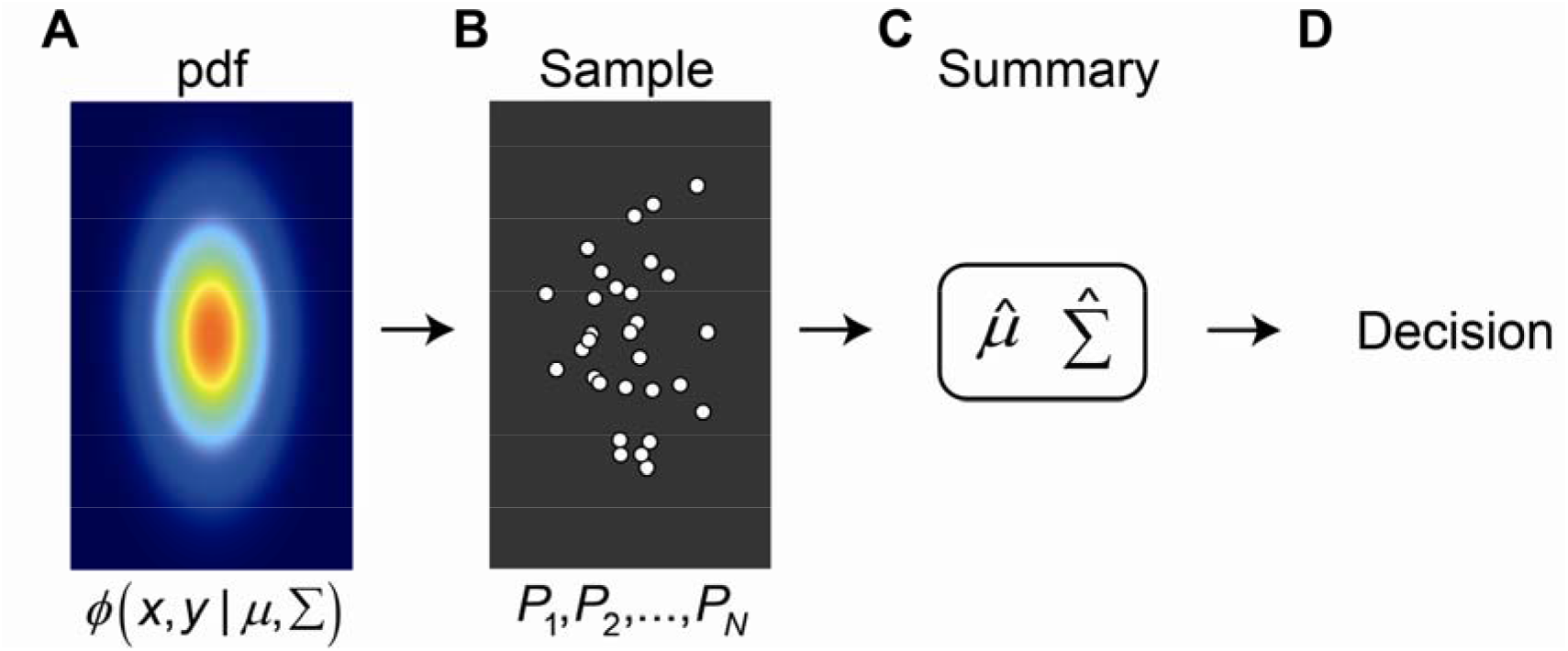
Parametric decision making based on a sample. **A**. Bivariate Gaussian PDF (referred to as the “population”). The population PDF is not known to the decision maker. **B**. The decision maker is given only a sample ***P*_1_, …, *P_N_*** of size ***N*** drawn from the Gaussian. **C**. The Gaussian parametric decision maker reduces a large number of sample values to the values of a small number of parameters. For the bivariate Gaussian, the sample is often reduced to five parameters that are estimates of population parameters 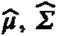 (referred to as “statistics”). **D**. The normative decision maker then makes decisions based only on these statistics, ignoring any “accidental” structure in the sample not captured by the parameters.

### Non-parametric approaches

At one extreme the decision maker could retain the entire sample in every detail. The summary is then the sample. If the sample size were large enough then the decision maker could use normative non-parametric approaches such as resampling (Efron & Tibshirani, 1993) that assume nothing about the population pdf. For example, he could estimate Eq. [1] by simply calculating the proportion of sample points that fall within the region ***T***. As sample size increases, this value converges in probability to the correct answer as a consequence of the weak law of large numbers (Wasserman, 2004). Here, though, we will work only with small sample sizes (5 or 30 points) where approaches based on resampling would lead to implausible predictions of human behavior.^3^ We consider instead parametric approaches (Lehmann, 1983).

### Parametric approaches

In parametric estimation we restrict the choice of pdf’s to a family of pdf’s summarized by a small number of parameter values corresponding to the selected pdf. Intuitively, the parametric decision maker “knows” more than just the sample. In Figure 2AB he knows that the pdf is unimodal and symmetric about two axes. For the bivariate Gaussian, in particular, the decision maker could replace the sample of size ***N*** shown in Figure 2B by the *summary* in Figure 2C comprising the sample mean 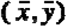, the sample covariance 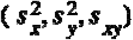 and the sample size ***N***. When the underlying pdf is Gaussian, these sample statistics are the corrected maximum likelihood estimates of population mean and covariance 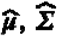. The estimates 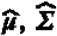 are *jointly sufficient statistics* (Berger, 1985; Hogg & Craig, 2002) that capture all of the data relevant to estimating the parameters of the population pdf. Not every pdf family has jointly sufficient statistics (Berger, 1985; Hogg & Craig, 2002).

Any invertible transformation of the estimates 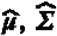 are also *jointly sufficient statistics* (Berger, 1985; Hogg & Craig, 2002) and replacing 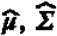 by the transformed parameters would lead to the same conclusions as those we reach. We refer to the model decision maker in Figure 2 applied to any judgment or estimation as the *normative BDT* decision maker for that task (the “normative decision maker” or “normative model” when context permits). We emphasize that this normative model is representative of a broad class of equally normative models based on jointly sufficienct statistics.

Summarizing the sample by 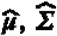 reduces a potentially large number of sample values to just five numbers, a remarkable degree of data compression, made possible by the parametric assumption that the sample is drawn from a bivariate Gaussian distribution. The decision maker can base his decision on this summary with no loss.

There are four advantages to basing all decisions in any task on a small number of parameters. The first, most obvious, is *parsimony*. The full complexity of the sample is replaced by a handful of estimated parameters. Second, the calculation of parameters (Figure 2C) does not depend on the task (Figure 2D). We use the same summary for all tasks. The third advantage is that the summary ignores *accidental structure* in the sample that provides no useful information about the underlying population. In Figure 2B, for example, the cluster of five points at the bottom of the sample is an accident of sampling. If we sampled again, we are unlikely to encounter a similar cluster. The cluster is visually salient but it would be a mistake to give more weight (or less weight) to the points in these cluster simply because they form an apparent cluster. In particular, the Gaussian pdf is symmetric about its mean in the vertical direction and this symmetry implies that the illusory 5-point cluster would be as likely to appear near the top of the sample – reflected about a horizontal line – as at the bottom where it is. The last advantage is that knowledge of the form of the pdf can potentially improve any non-parametric decision or estimate based solely on a sample. The parametric decision maker knows more than just the sample. Knowing, for example, that a pdf is bimodal even without knowing the locations of the modes is one example. Knowledge of the evident vertical and horizontal symmetry of the bivariate Gaussian pdf in Figure 2B is potentially useful in making decisions.

We will compare human performance to the normative Gaussian BDT decision maker sketched in Figure 2A-D. To compare human performance to normative BDT in detail, we measure the *influence* of each point ***P*** in the sample on the decision maker’s choices and on the choices of the normative model. Historically, influence measures were extensively used in the theory of robust estimation in mathematical statistics (Hampel, 1974; Huber & Ronchetti, 2009). and in research concerning depth cue combination (Maloney & Landy, 1989; Landy et al., 1995). We sketch here how we measure influence experimentally. The details are to be found in the Methods section.

### Influence measures

In Figure 3A, a bivariate Gaussian sample is shown in white circles. The underlying population pdf is drawn as a heat map and lightly sketched as elliptical contours of equal probability density. The decision maker’s hypothetical task is to estimate the probability ***P*[*T*]** that an additional point drawn from the same pdf will be above the green line, a region marked by ***T***. We wish to evaluate how each point in the sample enters into his estimate.

**Figure 3.**
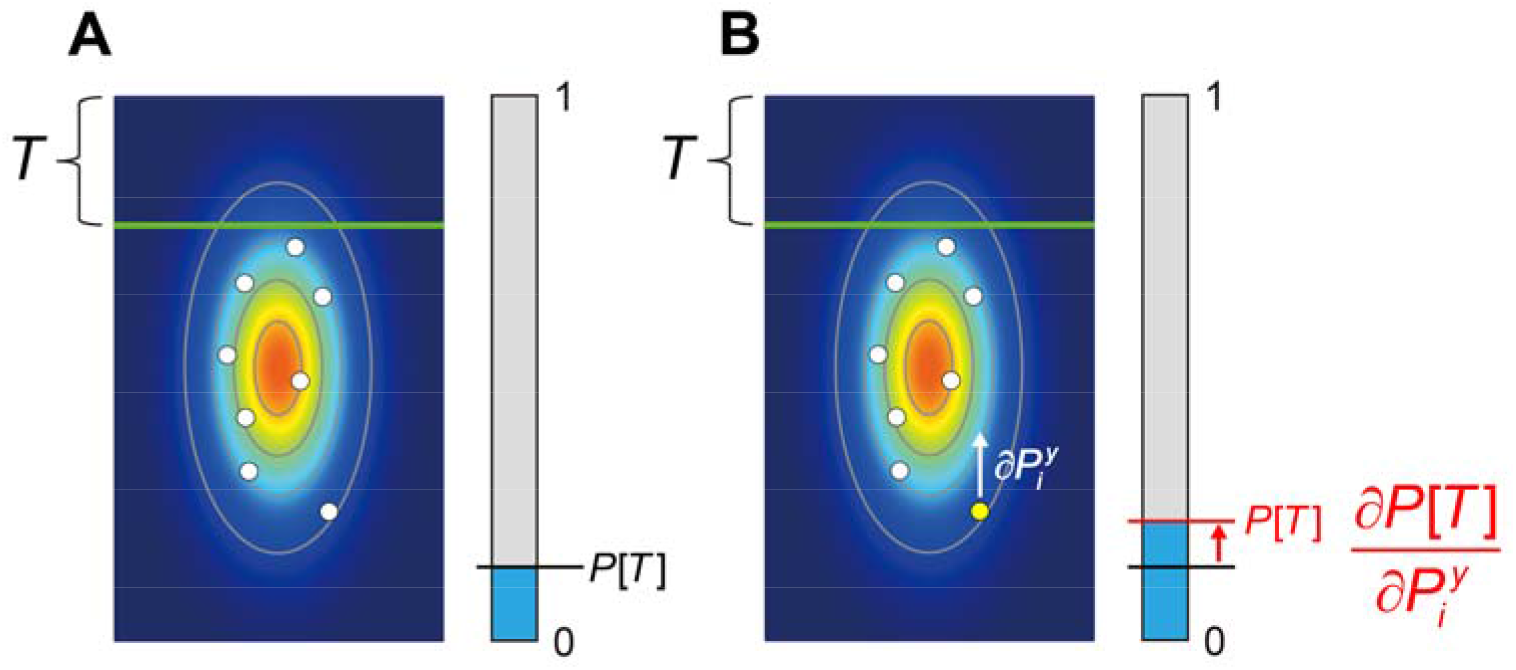
Measuring influence. **A**. A hypothetical experiment. A sample is drawn from a bivariate Gaussian pdf marked by a heat map and contours of equal probability density. The blue bar represents the decision maker’s estimate of the probability that an additional point drawn from the same underlying pdf will be in the region above the green line ***T***. The precise task is not important. **B**. Measuring influence. The (vertical) influence of one point in the sample can in principle be measured by perturbing it slightly in the vertical direction and measuring the effect of the perturbation on the decision maker’s estimate ***P*[*T*]**. The ratio of the change in estimate to the magnitude of perturbation is the *influence* of the point on the setting. We do not use this method (single point perturbation) but instead use a method based on linear regression. See *Methods*. The influence measures allow us to characterize how each point in the sample affects the decision-making.

The decision maker’s estimate of the probability ***P*[*T*]** is shown in a blue/grey scale in Figure 3A. Now suppose we alter the sample by shifting a point ***P*** slightly upward. In Figure 3B, the change is shown as a vertical red arrow and it is exaggerated in size to make it visible. The small change in ***P*** may result in a shift of the estimate and we are in effect computing a numerical estimate of the gradient of partial derivatives of probability estimate with respect to the vertical direction 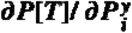, the influence in the vertical direction of the point. Intuitively, we could imagine grasping each point in turn and “wiggling” it up and down to see how ***P*[*T*]** changes. If the influence is 0, for example, then we would conclude that the point ***P*** is not used by the visual system in computing ***P*[*T*]**.^4^ We will estimate the decision maker’s ***empirical influence*** 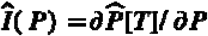 by regression analysis (see *Results*).

For any normative model we can also compute the ***normative influence I*(*P*) = *∂P*[*T*]/*∂P*** for any task we choose analytically, by numerical differentiation (Burden & Faires, 1985) or by Monte Carlo simulation. Influence is a signed measure and to compare empirical influence and normative influence we form the ratio

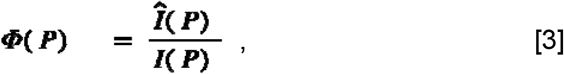

the ***influence ratio*** for any given point ***P***. If the decision makes use of the points as the normative model does, the influence ratios will all be 1. The influence ratio gives an indication of which points have too great or too little influence in absolute magnitude compared to normative. If the influence ratio is negative then the influence measure for the human decision maker is of opposite sign to that of the normative BDT observer. We could measure it for each of the five points in the cluster we discussed in Figure 2.

To summarize, in this article we test three properties of normative use of pdf’s derived from samples: ***accuracy*** (Eq. 1), ***additivity*** (Eq. 2), and ***influence***. We anticipate that participants will fail to match the normative BDT model perfectly. Our primary interest is in *patterned* deviations from normative. Such patterns might indicate what participants are actually doing in carrying out the task. Influence measures will allow us to seek such patterns (Maloney & Mamassian, 2009; Zhang et al., 2013, 2015).

## Results

### Testing accuracy and additivity

Participants first completed an interval estimation task designed to test both accuracy and additivity. Participants were shown a sample of 5 or 30 dots from an anisotropic bivariate Gaussian distribution (Figure 4A). The population means of the bivariate Gaussian distribution was fixed on a center of the screen, whereas the population covariance (the “size” and “shape” of the Gaussian pdf) randomly changed with each trial.

**Figure 4.**
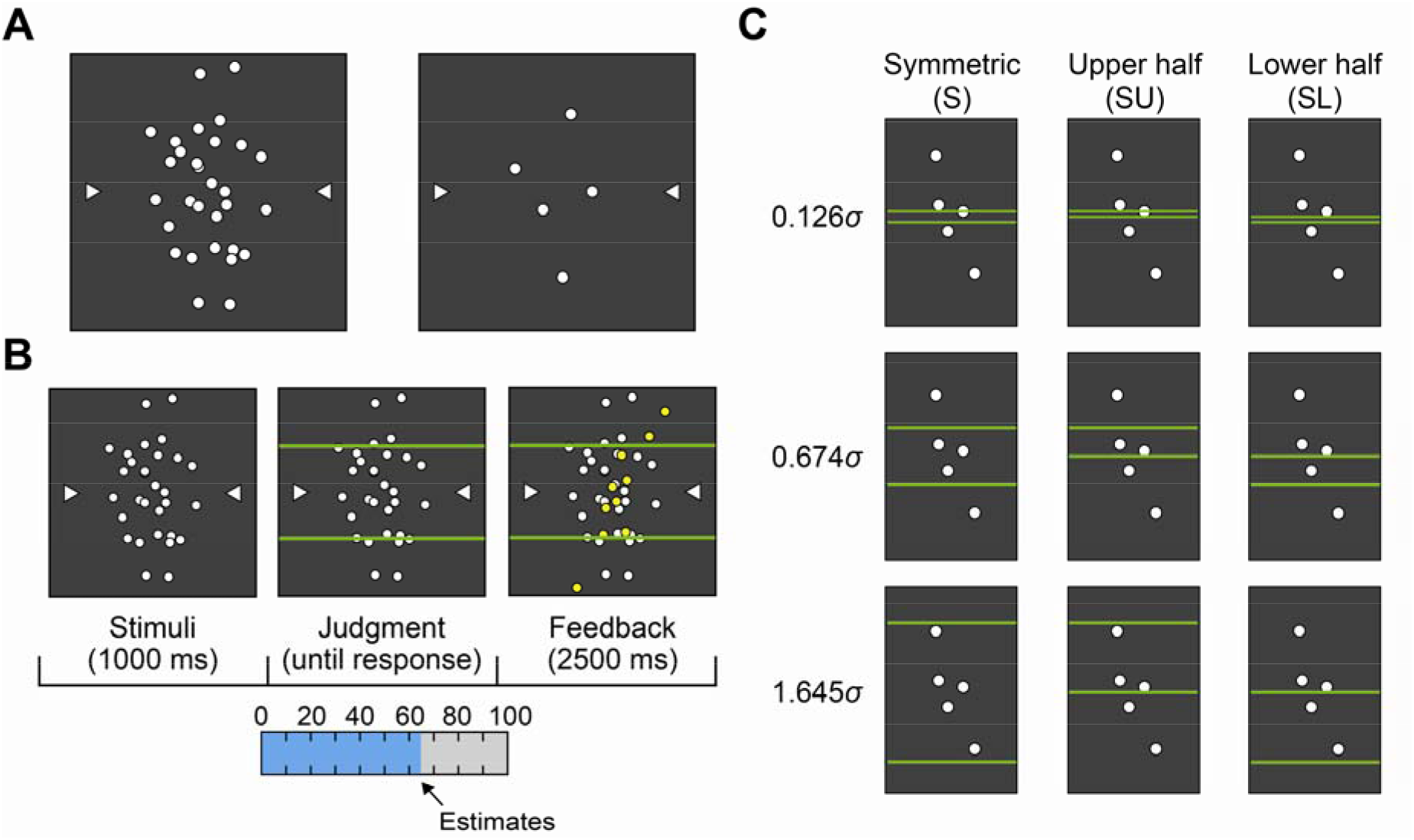
Design of the interval estimation task. **A**. Examples of 30-dot and 5-dot samples. The horizontal and vertical coordinates of the dots are independent random variables drawn from a bivariate Gaussian distribution. **B**. The trial sequence of the interval estimation task. The sample appears and then two horizontal lines. Participants judged the probability that an additional sample from the same distribution would fall into the region delimited by the horizontal lines. **C**. Three configurations of interval estimation with respect to the center of the screen. The upper half and lower half intervals are non-overlapping; their set-theoretic union is the symmetric interval. The vertical interval extents are expressed as in proportions of σ, the standard deviation of the population pdf in the vertical direction.

After a fixed interval (1 sec), two green horizontal lines across the distribution of the white dots appeared (Figure 4B). Participants were asked to judge the probability that a new dot drawn from the same population mean and population covariance would fall within the region delimited by the two lines. Accurate performance in this interval estimation task was equivalent to integrating the probability density of the.Gaussian distribution within the specified region (Eq. 1). We compared decision makers’ estimates to the correct estimates (a test of *accuracy*).

Participants recorded their probability from 0% to 100% by moving a digitized pen horizontally (Figure 4B). After their judgment, 10 new, yellow dots drawn from the same distribution that generated the sample were presented for 2.5 sec. The new dots fell inside or outside the horizontal lines giving participants performance feedback concerning their probability estimates.

The horizontal lines appeared in any of three configurations with respect to the center of the screen. The lines covered a symmetric interval ***S***, the upper half of the symmetric interval ***SU***, or the lower half of the symmetric interval ***SL*** (Figure 4C). The two half regions are non-overlapping and together form a symmetric region. These triples allowed a test of *additivity*: ***P*[*S*] = *P*[*SU*] + *P*[*SL*]**. We varied the interval width to make nine probability conditions equally spaced from 0.1 to 0.9 for the symmetric interval ***P*[*S*]** and from 0.05 to 0.45 for the asymmetric intervals ***P*[*SU*]** and ***P*[*SL*]**. Participants saw all three types of intervals on different trials. They repeated each condition 5 times, resulting in 270 (2 sample sizes × 3 configurations × 9 probabilities × 5 times) trials in total.

#### Test of accuracy

We tested whether the decision maker’s estimates of all intervals *S, SU, SL* were accurate. Before doing so, we confirmed that their estimates did not change across trials (Supplementary Figure 1). We then averaged the data over the five trials for each probability condition.

Figure 5A illustrates the mean estimates for a symmetric interval 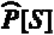 (averaged across participants and number of trials) against the correct probability ***P*[*S*]** of 30 dots and 5 dots, respectively. The observed deviations of the estimates for the symmetric interval is similar to the patterns of distortion in decision under risk (Gonzalez & Wu, 1999; Prelec, 1998; Tversky & Kahneman, 1992): participants systematically overestimated the probability induced by small and medium intervals.

**Figure 5.**
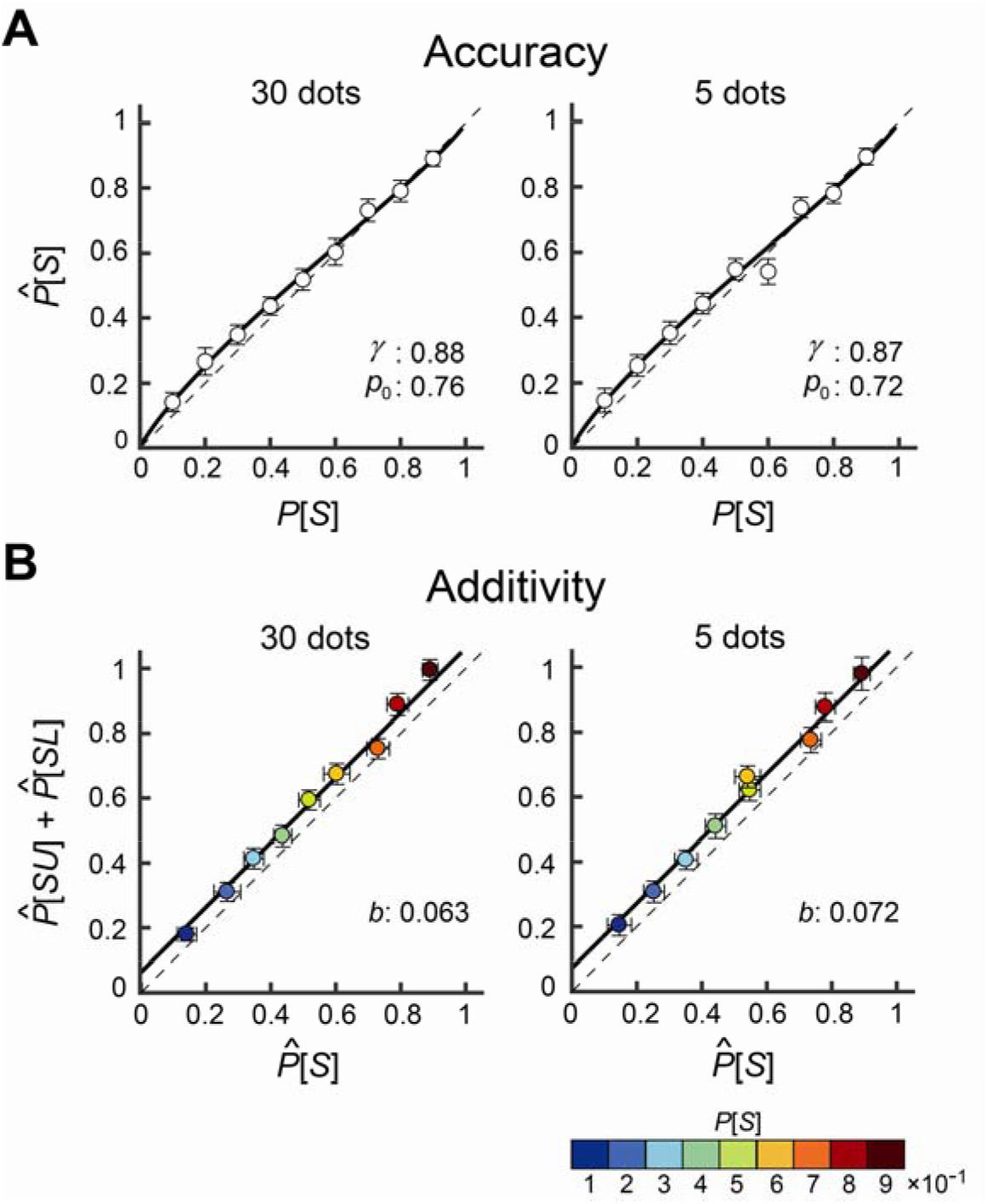
Results of the interval estimation task. **A**. Accuracy. The decision maker’s mean estimates of probability in the symmetric interval 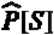 are plotted against the correct probabilities ***P*[*S*]**. The black thick curve is the maximum likelihood estimate of a linear-in-log-odds function fitted to the data. See text. **B**. Additivity. The sum of estimates in upper and lower halves of the symmetric interval 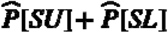 is plotted against the estimates of the symmetric interval 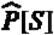. The color scale of the circle indicates the correct probability between 0.1 and 0.9. The black thick line is the best-fit estimate by a super additive function. For A & B, data is averaged across the decision makers, and the error bars indicate **±** 2 s.e.m.

There are several models used to model distortions in the estimates of probability (Gonzalez & Wu, 1999; Prelec, 1998; Tversky & Kahneman, 1992). Zhang and Maloney (2012) used linear transformations of the log odds of probability. The linear in log odds model (LLO) is defined^5^ by the implicit equation

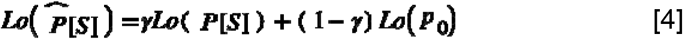

where 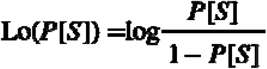 is the log odds (Barnard, 1949) or logit function (Berkson, 1944). Eq. [4] has two free parameters. The parameter ***γ*** is the slope of linear transformation of the log odds of the probability and ***p*_0_** is the “crossover point” where the probability distortion function crosses the identity line (see Zhang & Maloney (2012) for details).

We also fit other probability distortion functions taken from Tversky and Kahneman (1992) and Prelec (1998) and found that the LLO function provided consistently better fits to the data (Supplementary Table 1). See Luce (2000), Zhang & Maloney (2012) and Zhang et al (2020) for additional discussion of these models and their near equivalence.

We fit the LLO Model to the mean estimates across decision makers (Figure 5A) using maximum-likelihood estimation. The thick curves in Figure 5A shows the estimated LLO functions. The null hypothesis of no distortion is (in terms of the LLO parameters) ***γ* = 1** with ***p*_0_** set to any value. Supplementary Table 1 summarizes the results of AICc model comparisons. We found that the LLO function was 629 times (in the 30 dot condition) and 2.2 times (in the 5 dot condition) more likely than the null hypothesis of no distortion. In brief, there was considerable support for probability distortion in the 30 dot condition but not in the 5 dot condition.

The estimated parameter values (***γ*** = 0.88 and ***p*_0_** = 0.76 in the 30 dot condition, ***γ*** = 0.87 and ***p*_0_** = 0.72 in the 5 dot condition) indicate that the participants overestimated small probabilities. The probability weighting function found in the decision under risk literature typically has a value ***γ*** = 0.5~0.8 (Wu & Gonzalez, 1996) and the cross over point is found to be around 0.3~0.4, which results in a slightly more concave function below the cross over point (overestimating small probabilities) and more convex function above the cross over point (underestimating moderate to large probabilities). The estimated values of ***γ*** we find are only roughly consistent with the decision under risk literature; the estimated values of ***p*_0_** are markedly larger. The individual plots for the estimates for each decision maker are avaialble in Supplementary Figure 2.

We repeated the analysis for *SU* and *SL*, the two halves the of symmetric interval. Probabilities were overestimated (Supplementary Figure 3). The LLO model fit the estimates of *SU* and *SL* best and the recovered values of ***γ*** and ***p*_0_** were similar to those in the estimates of the symmetric interval (Supplementary Table 1). To summarize, we found distortion (“overestimation”) in the probability estimates based on the sample when the induced probabilitity was small to medium; the estimates were close to accurate for values of probability near 1. The pattern of distortion were similar regardless of the number of samples provided to the participants or whether the decision maker judged the full interval or one of the half intervals *SU* or *SL*.

#### Test of additivity

The decision makers overall misestimated the probability associated with target regions: we next test whether they can accurately sum these erroneous probabilities across disjoint target regions or whether they make an additional error, failing additivity. In Figure 5B, we plotted the sum of the mean estimates (across the decision makers and trials) in the two disjoint regions 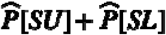 against the mean estimates in the single symmetric region 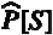. We know that these estimates are distorted but we wish to test whether the observer’s estimates of the sum of the observer’’s own estimates are systematically sub- or super-additive. The individual plot shows super-additivity for many decision makers (but not all; Supplementary Figure 4). We tested for failures of additivity consistent with the model:

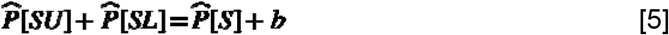

The null hypothesis of additivity is (in terms of the parameters) ***b* = 0**. A failure of additivity can be super-additive (***b* > 0**) or sub-additive (***b* < 0**).

We fit the three models to the mean estimates across decision makers (Figure 5B), and found that the super-additive model outperformed the other models (Supplementary Table 2). The super-additive model was 1660 times and 9336 times more likely than the null hypothesis of additivity for the 30 and 5 dot condition, respectively.

The thick curves in Figure 5B shows the super-additivity functions obtained by the fit for the 30 point and 5 point conditions On average, the sum of 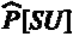 and 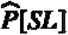 was 6.3% larger than the estimates of 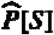 in the 30 dot condition and 7.2% in the 5 dot condition.

### Effects of probability distortions and super-additivity on movement planning tasks

The results showed that the participants have highly patterned failures in both tests of accuracy and additivity but these failures are small in magnitude. We might ask, how would the observed failures affect typical movement planning tasks? We examine how the pattern of probability distortion we find would have affected performance in analogues of two experimental tasks reported in the BDT literature.

In Figure 6A, we show a red penalty region with −100 points and a green reward region with +100 points from one of Trommershäuser et al. (2003b)’s experiments. Participants attempt 100 movements (black circles) toward an aim point (red diamond). We set the aim point shown in Figure 6A to maximize the expected reward of a typical participant given his motor variance. The objective probabilities of hitting within the red region, red and green region, green region, and outside of both regions are 0.002, 0.035, 0.879, and 0.084, respectively. The resulting expected gain is 87.31 points. If a hypothetical participant misestimates probabilities in a fashion consistent with our results, those probabilities would be transformed into 0.005, 0.058, 0.868, and 0.123, respectively, by the observed linear log-odds function in Figure 5A. An overestimation of the probability of hitting the red region along with an underestimation of the probability of hitting the green region reduces the expected gain to 85.64 points (a loss of 1.91%). The effect on the optimal aim point is slight. A green diamond shows the optimal aim point taking the observed LLO function into account and it almost completely overlaps with the red diamond. The failures of accuracy we observe would have essentially no effect on human performance.

**Figure 6.**
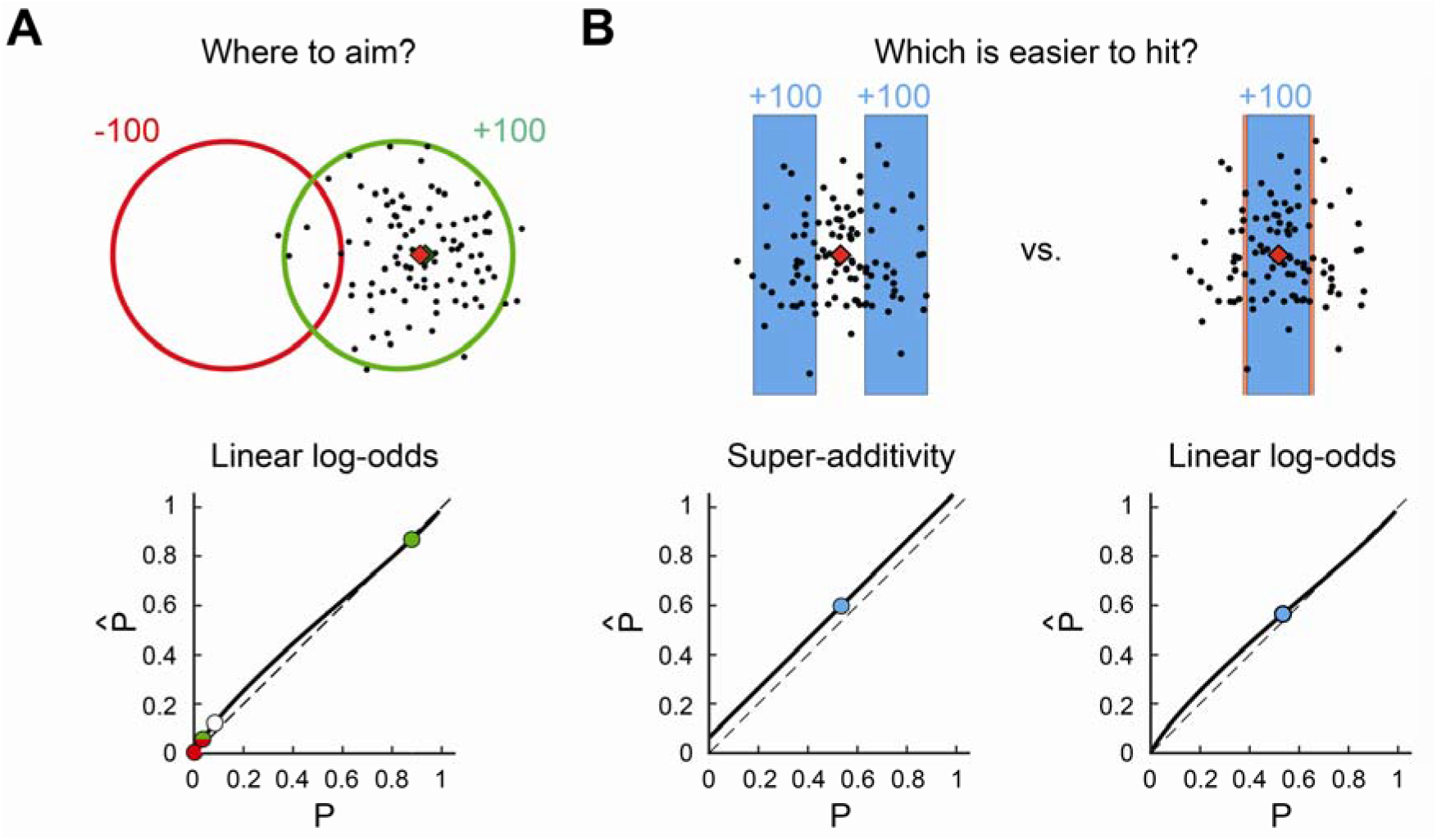
Hypothetical costs due to failures of accuracy and additivity in two previous experiments. **A.** A stimulus from Trommershäuser et al. (2003b). We chose the median stimulus from their experiment. In their task, the participant made speeded reaching movements to the reward region (green circle). The red circle denotes the penalty region. The distance between the two circles is 1.5 times of the radius of the circle. The radii of circles were 8.97 mm. A touch within the green region earns +100, within the red, −100, and within the green and red, 0. Hitting outside of both regions earns nothing. Black circles denote a possible isotropic bivariate Gaussian distribution of end points around the aim point (SD 3.89 mm, the average SD in Trommershäuser et al. 2003b). Given the standard deviation of the bivariate Gaussian distribution, the optimal aim point maximizing the expected reward was calculated and is shown as a red diamond. The objective probabilities of hitting each region with a possible end points are plotted against the subjective probabilities as circles. We consider a hypothetical participant who overestimate small probabilities by the linear-in-log-odds function in Figure 6A (30 points) with ***γ* = 0.88, *p*_0_ = 0.76**. The probability distortion slightly shifts the optimal aim point (with the new aim point shown as a green diamond almost completely covered by the red diamond). **B.** A stimulus similar to stimuli in Experiment 1 in Zhang et al. (2015). They used a two-alternative forced choice task. One of the options was a large, single rectangle target and the other comprised three disjoint smaller rectangles. To simplify our example, we replace the triple target by a double target. Hitting in either colored bar of the double target earned full reward. Participants decided which target (single or double) to attempt to hit and made speeded reaching movements to the center of the chosen target. Hitting within the rectangle earned the same reward. We used two target rectangles for simplicity. The standard deviation of the reaching movement was chosen to be 3.05 mm (the average of the participants’ measured SD’s in experiment 1 in Zhang et al. (2015)). The widths of the two rectangles are 1.5 times the SD and the gap between two rectangles is 0.75 times of the width of that rectangle. These widths and gap correspond to a median values of the targets used in Zhang et al 2015. The heights of the rectangles are set so that virtual participant’s end points do not fall outside the vertical boundaries. The width of the single rectangle is adjusted so that the objective probability of hitting the single target is that same as that of hitting the double target. The normative decision maker would pick each target 50% of the time. As a consequence of distortion of probability and super-additive, the decision maker instead picks the double target more often. If the single target is slightly increased in width by 14.7% (shown in a light red border), the decision maker would pick them equally often though his chances of hitting the single target are objectively greater.

In Figure 6B, we consider a variant on a task used by Zhang et al. (2015) in their experiment 1. Participants were asked to complete a two-alternative, forced choice task where one of the target options comprised three disjoint smaller rectangles (hitting anywhere in any of the three rectangle earned the reward) and the other was a larger single rectangle. We simplify their task to have two equally-sized disjoint targets (Figure 6B; left). The objective chance of hitting the double target is 0.534. We scale the width of a larger single target so that the objective chances becomes equal. If a hypothetical participant is super-additive, the probability of hitting the double target would inflate to 0.597 (Figure 6B; right). The estimates for the single target is 0.564 based on the LLO probability distortion function. While the normative BDT decision maker picks the double and single targets equally often, the decision maker with failures of accuracy and additivity will pick the double target more often than normative. The single target would have to increased in width by 14.7% (from 4.453 to 5.107 mm; this increase can be seen in a light red border) to restore indifference between the targets. The failures we find in the test of accuracy and additivity are small but patterned but have only modest effect on expected value in the tasks considered.

### Measuring influence

We designed a separate task to measure influence, the decision task. As in the interval estimation task, a sample of white dots (N = 5 or 30) is drawn from an anisotropic bivariate Gaussian distribution (Figure 7). The population covariance randomly changed with each trial. In this task, the participants could rigidly move (translate) the visible sample to any location on the screen.

**Figure 7.**
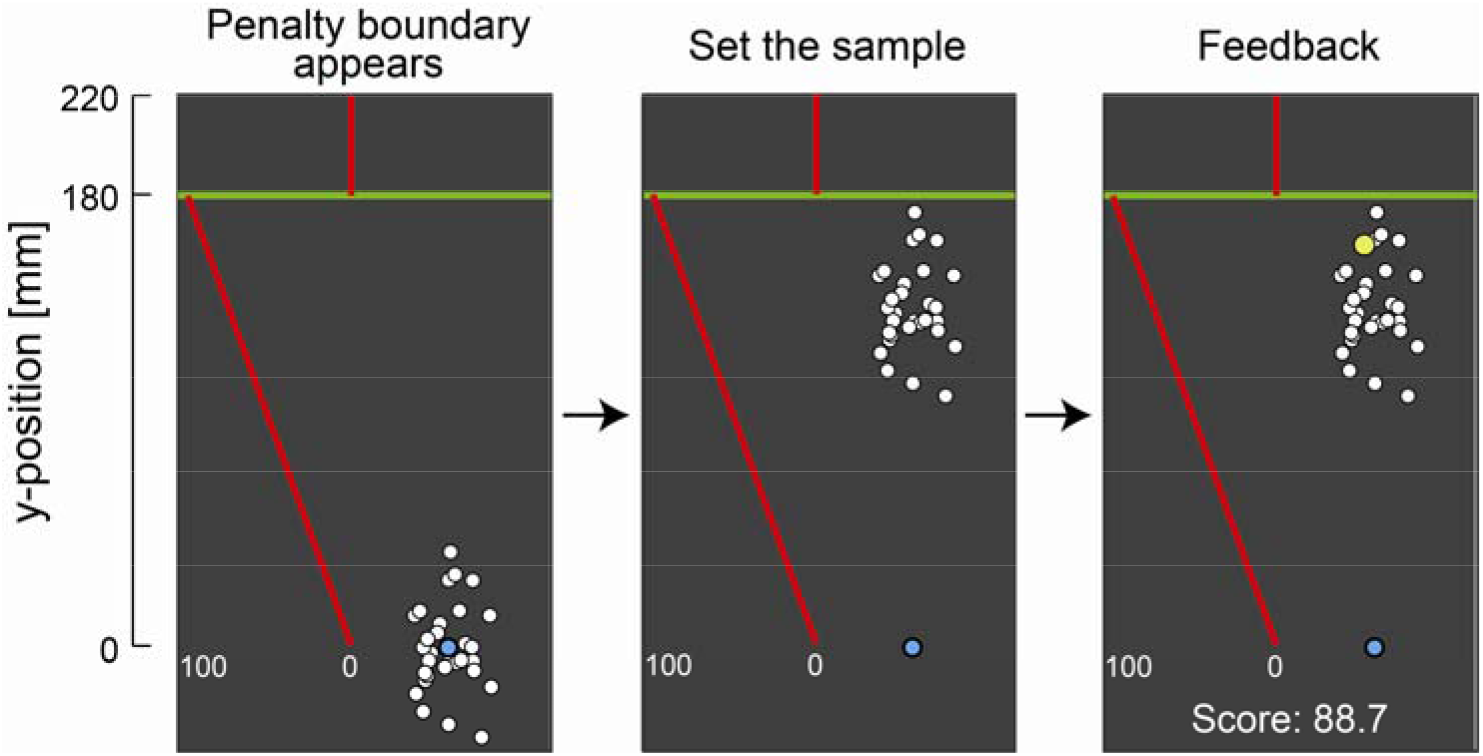
Decision task. In each trial, a sample of 30 or 5 dots was drawn from an invisible bivariate Gaussian distribution and shown on a visual display. The decision makers were asked to move the sample up and down from the blue starting point. In moving the sample the decision maker also moved both the sample and the pdf of the underlying invisible distribution. A penalty and 500. The red lines on the screen signaled the penalty and reward condiitons on each trial. After the decision maker set the location of the sample and its underlying pdf, one yellow dot was drawn from the same distribution. There were two penalty conditions, 0 and 500. The red lines on the screen signaled the penalty and reward conditions on each trial. If the yellow dot appeared above the green line, the decision maker incurred the penalty. If the yellow dot fell on or below the green line, the decision maker received a reward that was inversely related to the distance between the yellow dot and the green line. The rewards ranged from 0 to 100 points. In the figure the decision maker receives 88.7 points. The decision maker had to trade off the increased probability of a penalty if he moved the sample upwards and a reduction in reward if he moved it downwards.

The participants first set the sample to an initial position (blue circle) to start the trial. Next a green penalty boundary was shown (180 mm above the start). The participants then decided where to set the sample on the screen (Figure 7). We recorded the final vertical coordinate of the digitized pen as the participant’s set point in the trial. The sample could be moved to the left or the right but such horizontal movement had no effect on the reward or penalty incurred.

After the participants had chosen a location for the sample, a new point would be drawn from the pdf (now at the location set by the participants). The location of the new point with respect to the penalty boundary would determine the score in the trial. If the new dot was below the penalty boundary, the participants received a reward that was a decreasing linear function of the distance to the penalty boundary. However, if the new dot appeared above the penalty boundary, they incurred a penalty, accompanied by an aversive sound. There were two penalty conditions 0 and −500. Red lines on the screen denoted the reward functions.

The participant’s task was to maximize the total score by deciding where they locate the sample. Intuitively, if they left it far below the penalty boundary, the new dot would tend also to be far below the penalty boundary. They would rarely incur a penalty but any reward they received would tend to be small. If they moved the sample closer to the penalty boundary, then the probability of a penalty increased but the reward would tend to be larger. The decision maker in effect had to trade off between the magnitude of reward and the probability of penalty.

The design was factorial: two sample sizes crossed with two values of penalty. The trials from the resulting four conditions were interleaved and participants completed 50 trials in each condition for a total of 200 trials.

#### Calculating the optimal set point

The decision maker is given only the sample data from the population pdf and does not know the population pdf or its parameters. Thus the normative decision maker obtains the estimates 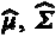 based on the observation of sample mean 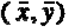 and sample covariance 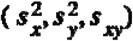 and he chooses a setpoint to maximize expected reward. But beware: the decision maker cannot treat the estimated parameters as if they were the true population parameters. The normative decision maker must take into account the uncertainty in the estimates 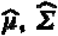 and allow for the difference in number of points in the 5-point and 30-point samples. See *Normative BDT model* in the Methods for how to maximize the expected reward given 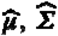 and the number of points.

There were no trends in decision maker’s set point from the beginning to the end of the task (Supplementary Figure 5). Therefore we computed the average set point across trials (Supplementary Figure 6A). A two-way within-participant ANOVA showed a main effect of the penalty condition (*F* [1, 16] = 42.68, *p* = 0.001, = 0.52) and a main effect of the number of dots (*F* [1, 16] = 53.08, *p* = 0.001, = 0.20). There was no significant interaction (*F* [1, 16] = 0.99, *p* = 0.34, = 0.001). This suggests that participants made a riskier decision in the 5 dots conditions compared with the 30 dots conditions and made a safer decision in the large penalty conditions compared with the small penalty conditions. To further compare human performance to the normative BDT, we analyzed how each point in the sample influenced the decision maker’s set point relative to the normative set point.

The normative influence of each point on the normative set point ***I*(*P*) = *∂S*/*∂P*** was computed numerically. The influence of each point on the decision maker’s set point 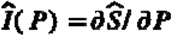 was measured by regression analysis (See *Measuring influence* in Methods). Figure 8A shows the normative influence and measured influence for each penalty condition and sample size. The sample points are sorted and assigned an order index ranging from 1 (lowest sample point) to either 5 or 30 (highest sample point) depending on sample size. The normative influence is skew-symmetric: higher points nearer the target region are assigned negative influences large in magnitude while sample points furthest from the target region are also assigned influences large in magnitude but opposite in sign. Measured influence in contrast is largest in magnitude for sample points near the target region but points far from the target region have almost negligeable influence. It might seem plausible that points far from the target region should be assigned little influence but the normative BDT decision maker does not do so. For the normative decision maker, the point nearest the target region and the point furthest from the target region have the largest influences (in magnitude) though opposite in sign.

**Figure 8.**
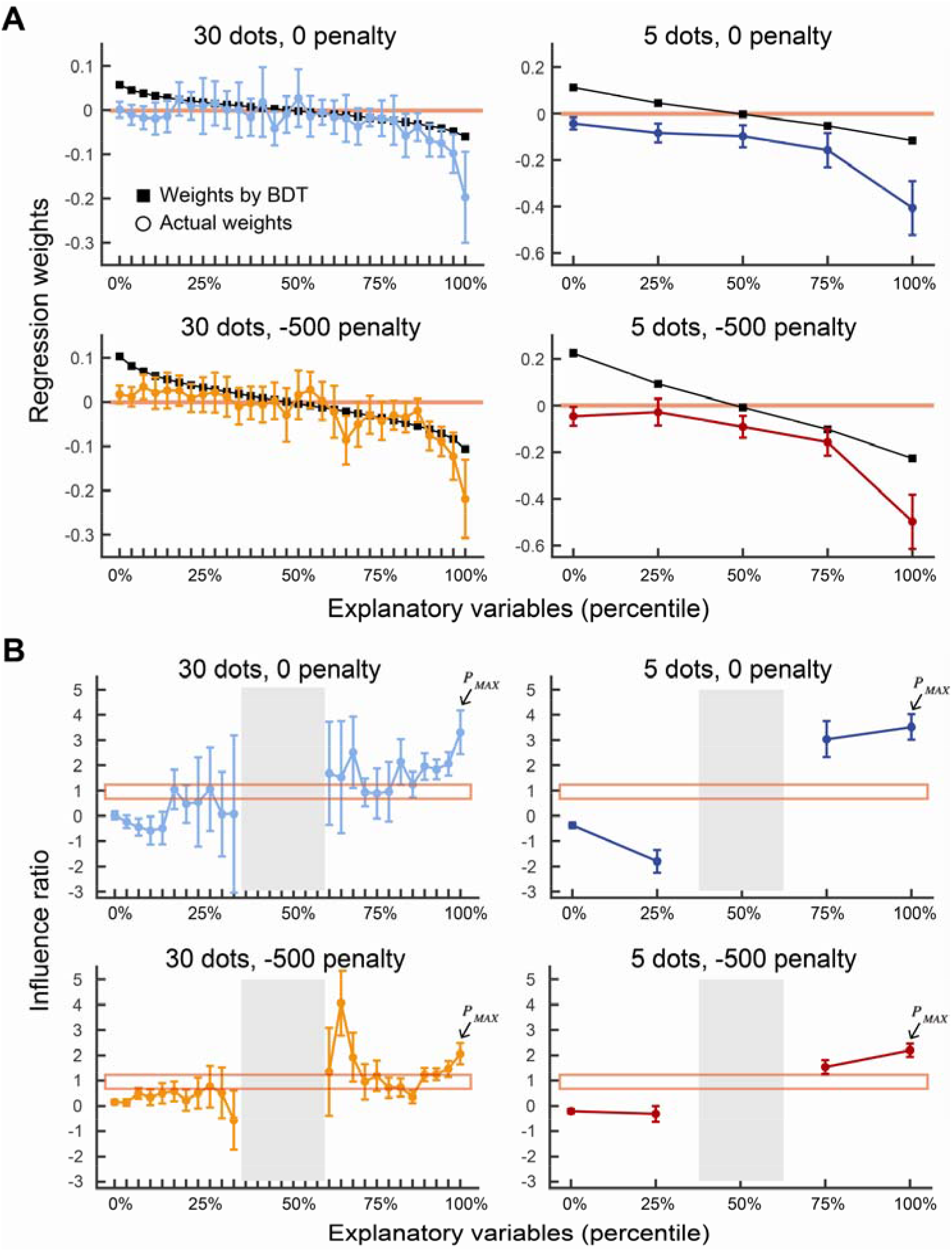
Measured influence and the influence ratio. **A**. Normative BDT influence and measured influence. We order the sample points for each sample point from 1 (the lowest) to either 5 or 30 (the highest) depending on sample size. We plot the the mean across observers of the estimated influence of each sample point on the participant’s actual set point versus its order index (colored circles). We plot the influence expected for the normative BDT decision-maker versus order index (black squares). Data is averaged across the participants, and the error bars indicate **±** 2 s.e.m. Negative influence indicates that a set point is set further away from a penalty boundary when a sample point is generated close to a penalty boundary relative to a starting point. The influence measures for the normative BDT model are skew-symmetric and a thin red line marks the axis of symmetry. The highest points (near the target region) and the lowest points (farthest from the target region) have influence equal and oppoitie in sign. The middle points have less influence. In contrast the human decision maker has influence measures that roughly decrease in magnitude as we go away from the target region. The lowest points in the sample have little or no influence. Sample points distant from the target region have little influence. **B**. Influence ratios. The average across observers of influence ratios (measured influence divided by normative BDT influence) for each sample point are plotted versus the order index of the sample point. A value of 1 indicates that measured influence was identical to the normative BDT influence Error bars denote **±** 1 s.e.m. The influence ratios deviate from 1 (makred by a thin red box). They are not. For the sample points nearest the target region the influence ratios are too large but they approach 0 for sample points far from the target region. The gray-shaded central range could not reliably be estimated due to the denominator (i.e., normative BDT influence) being near zero.

Figure 8B shows the corresponding influence ratios 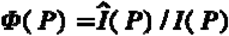. The sample points are again sorted and assigned an order index. We could not reliably estimate the ratio near the median point because the normative influence measures (denominator) there were almost zero (Figure 8A) and we omit them. The normative decision would have values of normalized influence consistently near 1 with no patterned deviations. Instead, there is a trend: the points close to the maximum point have equal or greater influence compared to normative whereas the points close to the minimum point have almost no influence or negative influence (the influence is in the opposite direction from normative influence). In particular, the influence of the maximum point is two to three times greater than normative (marked by arrows). The pattern of measured influence deviates markedly from the expected pattern derived from the normative BDT decision maker.

#### An alternative heuristic strategy

The results of the influence analysis suggest that participants gave considerable (and inapproriate) influence to the points closest to the target region above the green line. We consider an alternative heuristic strategy (the *max-point strategy*) where the participants set the ***P_MAX_*** point to a reference boundary internalized in their decision system (Figure 9A). The reference boundary could be anywhere on the screen and is a free parameter for the model fit. In the models, the set point is determined by moving the ***P_MAX_*** point until it reaches to the reference boundary ***B*** as below ***S* = *B* − *P_MAX_*** where the location of ***P_MAX_*** refers to the the location when the decision makers set the sample at the starting position.

**Figure 9.**
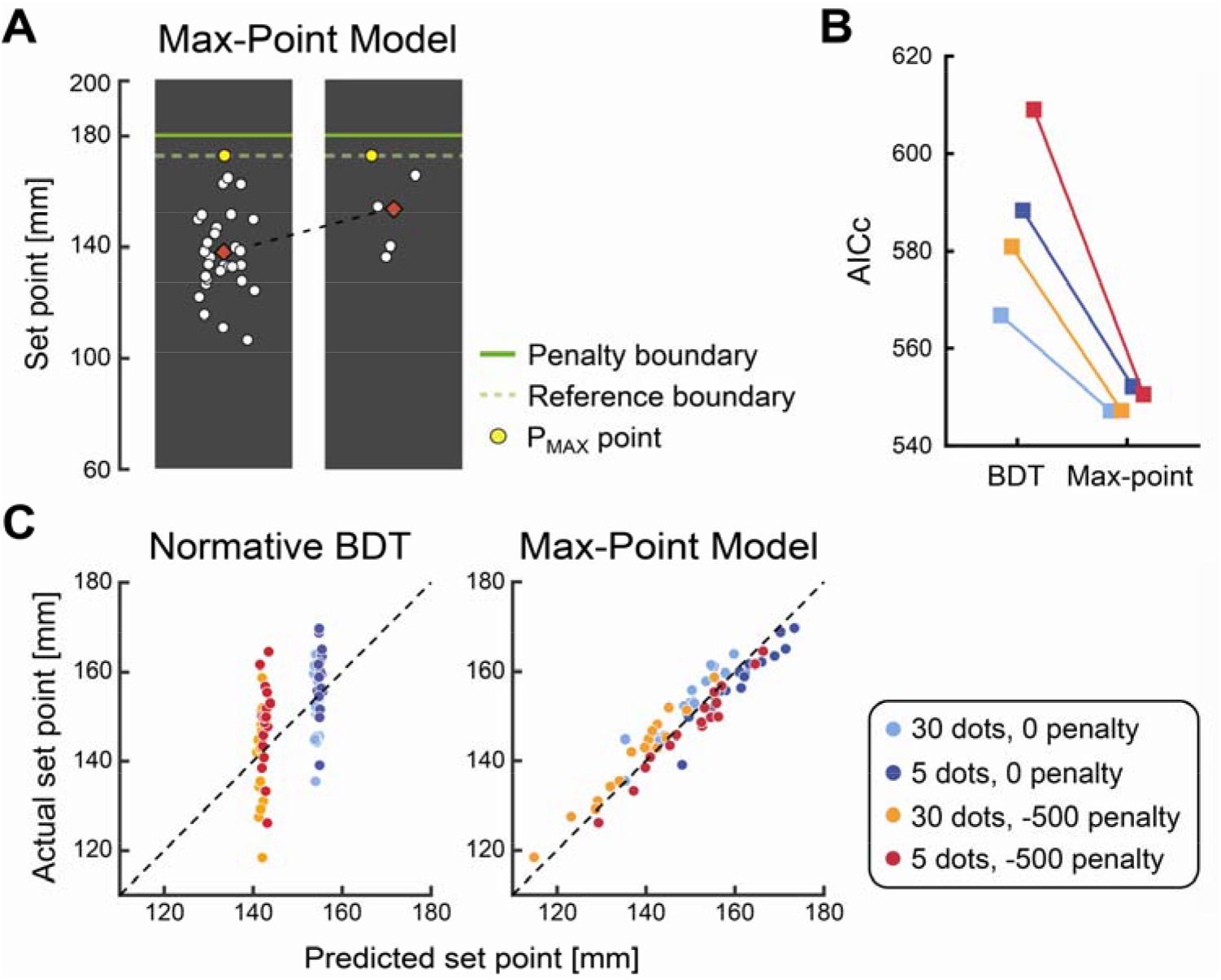
The Max-Point Model. **A**. Illustration of the Max-point model. Two samples drawn from the same pdf are shown, one of size 5 the other of size 30. The decision maker sets each sample so that the maximum point ***P_MAX_*** falls on a criterion boundary (dashed green line) chosen by the decision maker. The reference boundary is identical for 30-point samples and 5-point samples and as a consequence the mean settings (red diamonds) for the 5-point sample are markedly higher than those for the 30-point samples. **B**. An AICc model comparison of the normative BDT model and the Max-Point Model. The The lower the AICc, the better the model. **C**. A plot of mean settings for each participant verus the predictions of the normative BDT model. **D**. A plot of mean settings for each participant verus the predictions of the Max-point model. In sum, the Max-Point Model outperformed the normative BDT and reproduced the participant’s set point fairly well.

Therefore, the Max-Point model predicts a higher set point in the 5 dot than the 30 dot condition (Figure 9A). In Figure 9CD, we show correlation plots between the actual set point and the model prediction of normative BDT and the heuristic model. The normative BDT clearly failed to capture the behavioral pattern whereas the Max-Point model well matched. We calculated the AICc for each model, each participant, and each condition (Figure 9B). On average across conditions and participants, the Max-Point model was **1.2 × 10^12^** times likely than the normative BDT, suggesting that, in their decision, the decision makers primarily relied on the extreme point ***P_MAX_*** rather than the parametric estimates of the population pdf.

To summarize, the influence estimates we obtain from human decision makers deviate from normative. The sample points nearer to the green boundary are assigned much more influence that the normative Gaussian BDT model would assign. Points distant from the target region are assigned influences near 0. The decision makers are making their judgments on subset of the sample points near the target region, ignoring those further away. Their error was costly, leading to a larger number of penalty trials in the 5 dot than 30 dot condition (Supplementary Figure 6B). The actual total score was on average 82.7% (57.0% ~ 95.7%) relative to the BDT theoretical maximum score (Supplementary Figure 6C). Decision makers have an efficiency of 82.7%: their choices of strategy cost them 17.3% of their potential winnings.

## Discussion

Bayesian Decision Theory (BDT) is used to model ideal performance in a wide variety of experiment tasks in human perception, movement planning, and cognition (Ma, 2019; Maloney & Zhang, 2010; Rahnev & Denison, 2018). It captures important aspects of human performance in everyday tasks: the outcomes of these tasks entail rewards or penalties for the decision maker and the decision maker is only partially in control of the outcome. The decision maker can choose an action (make a decision) but the choice of action does not determine the outcome.

BDT combines information about uncertainty and rewards to maximize expected reward. Experimental tests of BDT often estimate the *efficiency* (Geisler, 1989; Trommershäuser et al, 2003ab) of the human decision maker – the ratio of human winnings to the maximum expected possible winnings predicted by BDT, arguably the key measure to consider in evaluating human performance. If efficiency is substantially less than 1 then we have evidence that the BDT model is not appropriate as a model of human performance. Past experimental tests of the BDT model in a wide variety of cognitive, perceptual and motor tasks have decidedly mixed outcomes, some finding that overall human performance appproaches the efficiency limit dictated by BDT while other find a marked gap between human performance and optimal (see Rahnev & Denison, 2018 for review).

Maloney & Mamassian (2009) and Zhang et al (2013) argue that a comparison of overall winnings to the maximum possible is not a powerful test of the hypothesis that BDT describes how human observers make decisions. Heuristic rules different from BDT can achieve efficiencies as close as we like to 1 (Maloney & Mamassian, 2009) or good overall performance may be due to an unwitting choice of task (Zhang et al, 2013). The brain may appear to be Bayesian (Knill & Pouget, 2004; Pouget et al., 2013) but in reality it is doing something else. What we want to know about is that “something else.”

We break down the Bayesian computation into elementary operations and test human ability to carry out three of these operations. We considered visual cognitive tasks where the human decision maker is given a sample from a bivariate Gaussian probability density function (pdf) and must use it normatively (Figure 2A-D). The transition from sample to pdf is a key step in the BDT computation because the use of the estimated pdf allows parametric decision-making based on a handful of estimated parameters and ignores any accidental structure in the sample. We first tested *accuracy*, the ability to correctly estimate the probability that an additional point from the specified pdf will fall into any specified region (Figure 4). This ability is essential to the BDT computation when different regions carry different penalties (Figure 6A). We then tested *additivity*, the ability to estimate the probability of landing in a region composed of multiple disjoint subregions (Figure 4). We found small but patterned failures of both accuracy and additivity (Figure 5) but argued that the costs of these observed failures in accuracy and additivity were minor or almost negligible in motor tasks used in the literature of BDT (Figure 6).

We last measured the *influence* of each point in the sample on the decision makers performance (Figure 7). The transformation from sample to the estimated pdf leads to skew-symmetric in influence: the highest point and the lowest point are assinged large influences in magnitute and opposite in sign (Figure 8A). In contrast, the individual’s use of sample information in making decisions deviated markedly from skew-symmetric. Although the brain may appear to be Bayesian (Knill & Pouget, 2004; Pouget et al., 2013) in some tasks, it is doing something else.

How can we understand human perfomance when it deviates from normative? One theory assumes that decision-makers have a different representation of the probability density function than a normative decision-maker has (for instance, “size” and “shape” of the Gaussian pdf) (Rahnev & Denison, 2018). Some studies indeed approximate decision maker’s internal model as a wider or skewed distribution than actual pdf (Acerbi et al., 2012; Stocker & Simoncelli, 2006; Zhang et al., 2013). This account has a strong assumption that decision-makers still extract parameters of the pdf from a sample (a key step in the BDT computation) and make decisions based on a parametric estimation (Figure 2A-D). Examination of the influence of individual points suggests that human decision-makers ignore the transform of sample to a handful of estimated parameters. The Max-Point model we developed does not assume such a parametric decision-making. Rather the model assumes a heuristic decision rule based on just one sample point closest to the target region. Nevertheless, the model led to a much better fit to human data than BDT (Figure 9).

The failures of accuracy and additivity do not invalidate the claim that normative BDT is a useful approximation to human performance and a reliable model of how human decision makers will behave in experiments. However, the discrepancy between measured influence and normative indicates that the human decision maker is using information differently than the normative observer and even ignoring sample points far from the target region that the normative observer assigns great weight to. Our finding is broadly consistent with the fact that the visual information is weighted differently by the order of stimulus presentation (Juni et al., 2012), reference stimulus (Li et al., 2017), or visual appearance (Spitzer et al., 2017). In this study however, we developed a method to measure how much *each piece* of information in the sample influenced the decision maker’s action. We showed the evident asymmetry in influence: decision-makers assign weight on outlying sample points close to the target region two or three times greater than the normative while they ignore sample points far from the target region. A heuristic model based on the maximum sample point is consistent with the asymmetry in influence we found. We reject the normative Gaussian BDT model we began with, even as an approximation to human behavior.

## Methods

### Participants

Seventeen participants (mean age 21.9, range 19–30, 5 males) participated in the experiments. Informed consent was given by each participant before the experiment. All participants were not aware of the hypothesis under test.. Participants received US$12 per hour plus a performance-related bonus.

### Apparatus

Stimuli were displayed on a vertical monitor (VPIXX, VIEWPIXX, 514 mm × 288 mm). The monitor resolution was 1920 × 1080 pixels with a 60-Hz refresh rate. Participants were seated at a viewing distance of 60 cm. A pen-tablet was set in front of the monitor (Wacom Intuos Pro Large, workspace: 311 × 216 mm). Participants manipulated a digitized pen to carry out the tasks. The horizontal-vertical coordinates of the digitized pen were recorded at 60-Hz. All stimuli were controlled using the Matlab Psychophysics Toolbox (Brainard, 1997; Pelli, 1997).

### Tests of accuracy and additivity: the estimation tasks

The tests of accuracy and additivity were carried out in the same session. In both tasks, the stimulus was a sample of white dots drawn from an anisotropic bivariate Gaussian distribution with mean 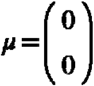 and covariance 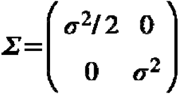. The origin was set at the vertical and horizontal center of the screen. On each trial, the population variance ***σ*^2^** was drawn from a uniform distribution on the interval 100 mm to 400 mm. The dots were round with radius 1.5 mm. There were two sample sizes as the number of white dots produced was N = 5 or 30. Two white triangles were presented at left and right sides of the area where the dots were displayed at the vertical mean ***μ_y_*** of the Gaussian pdf (Figure 4A). After presentation of the sample, two green horizontal lines (width: 1 mm) across the distribution of the white dots appeared (Figure 4B). Participants were asked to judge the probability that a new dot drawn from the same distribution would fall into the region delimited by the two lines. Participants recorded their probability by moving the digitized pen horizontally on a pen-tablet.

There were three configurations with respect to the center of the screen. The lines covered the range from **+ w*σ*** to **− w*σ*** in the symmetric interval ***S***, from the center of the screen to **+ w*σ*** in the upper half of symmetric interval ***SU***, and from the center of the screen to **− w*σ*** in the lower half of the symmetric interval ***SL***, where ***w*** means the interval width and ***σ*** means the population standard deviation. We varied the interval width to make nine probability conditions, [0.126***σ***, 0.253***σ***, 0.385***σ***, 0.524***σ***, 0.674***σ***, 0.842***σ***, 1.036***σ***, 1.282***σ***, 1.645***σ***]. The value of ***P*[*S*]** spanned the range 0.1 and 0.9 and ***P*[*SU*]** and ***P*[*SL*]** spanned the range 0.05 and 0.45. See Figure 4C.

There were 54 = 2 x 3 x 9 conditions created by combining two sample sizes (5 or 30 dot), three configurations (***S***, ***SU***, or ***SL***), and nine probabilities. Participants repeated the task in each condition of the 54 conditions 5 times (270 trials in total). We divided the experimental session into six blocks of 45 trials each. In the first three blocks, we presented the 30-dot conditions, in the remaining three blocks we presented the 5-dot conditions. Within each block, the configuration of horizontal lines was fixed but its order was balanced across participants.

### The decision task

The measurements of influence were carried out after the interval estimation task. The same participants completed this task after completing the previous estimation task. A sample of white dots (N= 5 or 30) was drawn from an anisotropic bivariate Gaussian distribution with mean 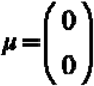 and covariance 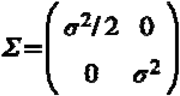. on each trial. The population variance ***σ*^2^** was drawn from a uniform distribution on the interval 100 mm to 400 mm.

In this task participants could move the sample rigidly to any place on the screen (Figure 7). To begin a trial, participants moved the sample to a blue initial position (radius: 4 mm). After holding for 1 sec, a white penalty boundary (PB) appeared 180 mm above the initial position. The PB turned green after a random interval (0.4-0.8 sec), which signaled the start of the trial. Participants moved the sample up or down deciding where to set the sample on the screen. They signaled their decision by pressing the button. After a button-press, a new yellow dot drawn from the same population pdf appeared and the vertical position of the yellow dot with respect to the green line determined the participant’s reward or penalty. If the yellow dot was above the green line then the participant incurred the penalty (0 or −500 points). If the position was on or beneath the green line by **0 ≤ *Δ* ≤ 180** millimeters then the participant received **(180 − *Δ*) / 1.8** points. That is, if the yellow dot were exactly on the green line (***Δ* = 0**) then the participant received 100 points. If it was 180 mm below the green line that participant received 0 points. The red lines in Figure 7 sketch the reward function and it was also shown on the screen as well as the score in the trial and the total score. The participants were instructed to maximize the total score and told that the obtained total score would convert into a bonus payment at the end of the experiment (75 cents per 1000 points).

There were four experimental conditions, two sample sizes (5 or 30 dot) and two penalties (0 or −500 points). Each condition was repeated for 50 trials, resulting in 200 trials in total. We divided the experimental session into 5 blocks of 40 trials. In each trial, either one of four conditions was randomly chosen. Participants received an average bonus of $9.70 (range $4.30 – $11.70).

### Normative BDT model

We modeled the normative setting point based on Bayesian Decision Theory (Berger, 1985; Maloney & Zhang, 2010). A data sample **(*P*_1_, *P*_2_, …, *P_N_*)** was randomly generated from the population pdf ***ϕ*(*x*, *y*|*μ*, *Σ*)**. A decision rule in the normative BDT is based on the estimated pdf 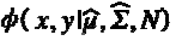. Although the population pdf is a bivariate Gaussian distribution, the horizontal coordinate is not relevant information as the reward is based on the vertical coordinate of new yellow dot. Therefore, we treated our model as a univariate pdf in the vertical dimension.

On each trial, the normative decision maker observes ***N*** data points in the sample **(*P*_1_, *P*_2_, …, *P_N_*)**. He converts the sample to the sample mean ***ȳ*** and sample variance ***s*^2^**, and utilizes these observations ***ȳ***, ***s*^2^**, and ***N*** to estimate the true population mean ***μ*** and population variance ***σ*^2^**. We denote the estimates of population mean and that of population variance as 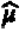 and 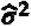, respectively.

To obtain those estimates, we assumed that the normative decision maker uses the knowledge of the generative model which produced the sample. In our experiment, we set the mean of the population pdf to a constant value. We can thus leave the estimates of population mean and focus on the estimates of population variance.

The prior probability of the estimated population variance can be written as a uniform probability density function:

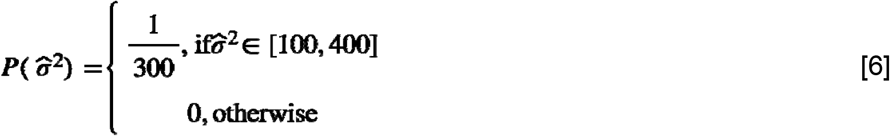

See Supplementary Figure 7A for this prior distribution.

The likelihood function of the estimated population variance depends on the number of points in the sample ***N*** and the sample variance ***s*^2^**. The estimates of population variance are distributed as chi-squared random variables with ***N* − 1** degrees of freedom and is described as a chi-square probability density function:

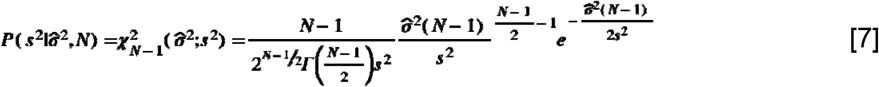

where ***Γ*( )** is the standard gamma function. Supplementary Figure 7B shows the example likelihood function when ***N* = 5** and ***s*^2^ = 200**. Supplementary Figure 8 also illustrates the examples with varing ***N*** and ***s*^2^**.

The normative decision maker computes the the posterior probability of the estimated population variance using a Bayes rule.

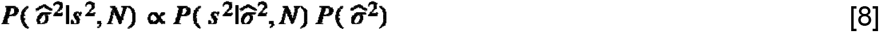

The posterior is proportional to the product of the likelihood and the prior. Supplementary Figure 7C shows the posterior probability distribution.

Once a particular population variance is estimated, we can determine the probability density function for producing the sample. Since the sample of 5 or 30 dot was randomly drawn from a bivariate Gaussian distribution, we set the estimated pdf in the form of a Gaussian distribution

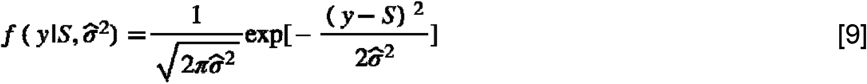

where ***y*** is a point in the vertical coordinate and ***S*** denotes a selected setpoint which shifts the center of estimated pdf. Supplementary Figure 7D illustrates three examples of the estimated pdf with varying 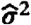 and fixed ***S***. Supplementary Figure 7E also illustrates the products of example pdfs with the reward function.

Given a setpoint ***S***, we can compute the expected reward by integrating the reward function ***G*(*y*)** and the estimated pdf 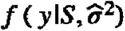 for each possible estimated population variance (Suppl. Figure 7F). We need to scale this expected reward by the posterior probability of the estimated population variance since the choice of 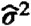 for the estimated pdf 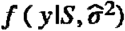 depends on 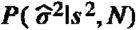. Supplementary Figure 7G shows the scaled expected reward as a function of 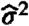.

The final expected reward can be obtained by integrating the scaled expected reward over the estimated population variance (Suppl. Figure 7H) and is defined as below.

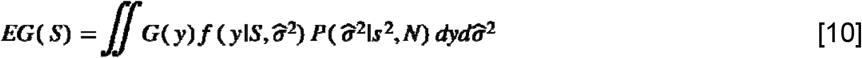

We changed the set point such that the expected reward can be maximized:

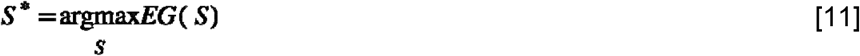

When ***P*_1_, , *P_N_*** are i.i.d. sample drawn from a normal distribution ***N*(0, *σ*^2^)**, the random variable 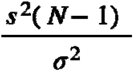 is distributed according to the chi-square distribution with ***N* − 1** degrees of freedom. Therefore, the probability of observing the sample variance ***s*^2^** given the population variance ***σ*^2^** can be written as:

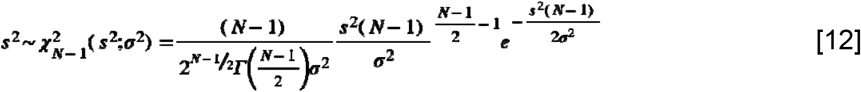

We can thus obtain the likelihood function of the estimated population variance in Eq. (9) by flipping ***s*^2^** and 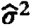 around.

### Measuring influence

#### Influence of each point on the human decision maker

We performed a regression analysis to estimate the influence of each point in the sample on the decision maker’s set point in the decision task. We ordered points in each sample (N = 30 or 5) in ascending order from the lowest sample point to 5 or 30 (highest sample point) depending on sample size (Figure 8). We used these explanatory variables to predict the final set point. We regressed the vector of decision maker’s set point **S** by the matrix of explanatory variables **X** containing a set of vectors from **P_MIN_** to **P_MAX_** to estimate the weight vector **W**. Our regression model is thus simply **S = X · W^T^**.

There are two issues if we used an ordinary least squares regression in this analysis. First, in the 30 dot condition, the number of explanatory variables (i.e., 30 variables) is relatively large to the number of observations (i.e., 50 trials). Under such a condition, the ordinary least squares regression tends to grossly overfit the training data and do not generalize the model prediction to the unobserved new data (Yarkoni & Westfall, 2017). Second, because the explanatory variables are order statistics, some variables are correlated with each other (e.g., the second maximum point possibly takes a large value when the maximum point is large whereas the second maximum point should be small when the maximum poiont is small). The regression weights can become poorly determined when there are many correlated variables in a linear regression model (Hastie et al., 2009).

We therefore used a ridge regression (L2-norm regularization) to estimate the weight vector **W**. Ridge regression is a machine learning technique which alleviate the problem of multicollinearity by adding the regularization term to the cost function (Hastie et al., 2009) as follow

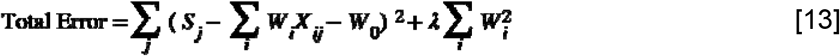

where ***S_j_*** is a set point at ***j***th trial, ***X_ij_*** is a ***i***th explanatory variable at ***j***th trial, ***W_i_*** is a regression coefficient for ***i***th variable, ***W*_0_** is the coefficient for a constant term ***λ***, and is a regularization parameter. In the ordinary least squares regression, the cost function to be minimized is solely the first term which is the sum of the squared difference between the data and model prediction. Ridge regression involves the second penalty term which is the sum of squared weight multiplied by regularization parameter ***λ***. Ridge regression basically shrinks the magnitude of the regression weights by increasing the amount of ***λ***. A large ***λ*** forces the regression weights to be closer to zero whereas a small has ***λ*** the opposite effect and in the limit as ***λ* → 0** the estimated weights converge to ordinary least squares regression weights.

To avoid overfitting, we performed 10-fold cross validation (Hastie et al., 2009; Yarkoni & Westfall, 2017). Specifically, for each decision maker and condition, we split the 50 data set to 10 equal-sized parts. Nine of ten parts (45 trials) were used for training data set and the remaining part (5 trials) were used for validation set. Given a particular ***λ***, we trained our model on training data set and then used the trained model to predict the total error on the left-out validation set. We repeated this process 10 times using different training and validation sets and we calculated the average error and standard error across 10 folds for each ***λ***. We used the “one standard error rule” (Hastie et al., 2009) for the choice of ***λ*** rather than the best ***λ*** with the minimized error, which results in selecting a more regularized model and preventing overfitting. The resulting estimates of regression weights are shown in Figure 8A.

### Influence of each point on the normative decision-maker

We computed the influence on the normative set point as follows. We assume the normative set point ***S*\*** as being a linear combination of weights and sample points 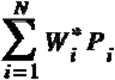. We induced a small change to a point ***P_i_*** while keeping the other points same. If there is a non-zero weight on ***P_i_***, a setting should change corresponding to the changes in ***P_i_***; otherwise the weight indicates zero. We denote the shifts in the normative set point with respect to the changes in a point as 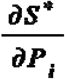 (i.e., a partial derivative). Since any other points are unchanged, the small changes in a particular point, ***∂P_i_***, is the only contributer to shift the set point and its amount depends on the amount of weight 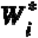. Therefore 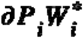 is a measure of the amount of the shifts with respect to that point (i.e., 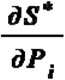) and we can derive the weight 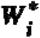 by 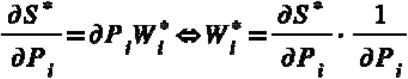.

We performed a Monte Carlo simulation to calculate the normative weights. We first generated a Gaussian sample of 30 or 5 points and computed the normative set point given a statistics of sample. In each sample, we induced changes in a point ***P_i_*** by 1 mm (due to the limitation of computational precision) and recorded the shifts in the normative set point given the changed statistics. We repeated the same process by generating a different sample 1,000 times and we defined the normative influence as 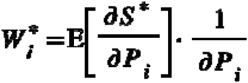. The simulated normative weights are shown in Supplementary Figure 8A.

## Competing interests

The authors declare no competing interests.

## Author’s contributions

KO and LTM conceived and designed the experiments. KO performed the experiments and analyzed the data. KO and LTM interpreted the results. KO wrote the manuscript and LTM revised the manuscript.

## Acknowledgments

We thank Jakob Phillips for his support in the experiment. This research was supported by Grant-Aid for JSPS Fellows No. 17J07822 awarded to KO and by a Guggenheim Research Prize awarded to LTM.

## Footnotes

1 See, for example, Trommershäuser et al. (2003ab; 2008). We assume that small shifts of aim point do not alter the covariance of the Gaussian distribution of end points. The argument presented here is easily modified to allow for changes in the covariance of the pdf with changes in aim point.

2 The probabilities for any target all together completely determine the pdf (Riesz-Fischer Theorem). We can in principle reconstruct the pdf from the target probabilities. Maloney & Mamassian (2009).

3 With a sample of size five and a target that is small compared to the interpoint spacings in the sample, for example, the only possible probabilities would be 0.0, 0.2, …, 1.0 and most would be 0.0 (no point within the target) and 0.1 (one point within the target).

4 We could similarly estimate the influence of the point in the horizontal direction but for the remainder of this article we focus on influence in the vertical direction only; for simplicity we will leave out the y-superscript in all following equations.

5 We write the equation in terms of the symmetric interval S for convenience. It applies equally to SU, SL as well.

## Notes

### Competing Interest Statement

The authors have declared no competing interest.

