## Supplemental information for "Using influence measures to test normative use of probability density information derived from a sample"

### 1 Supplementary results

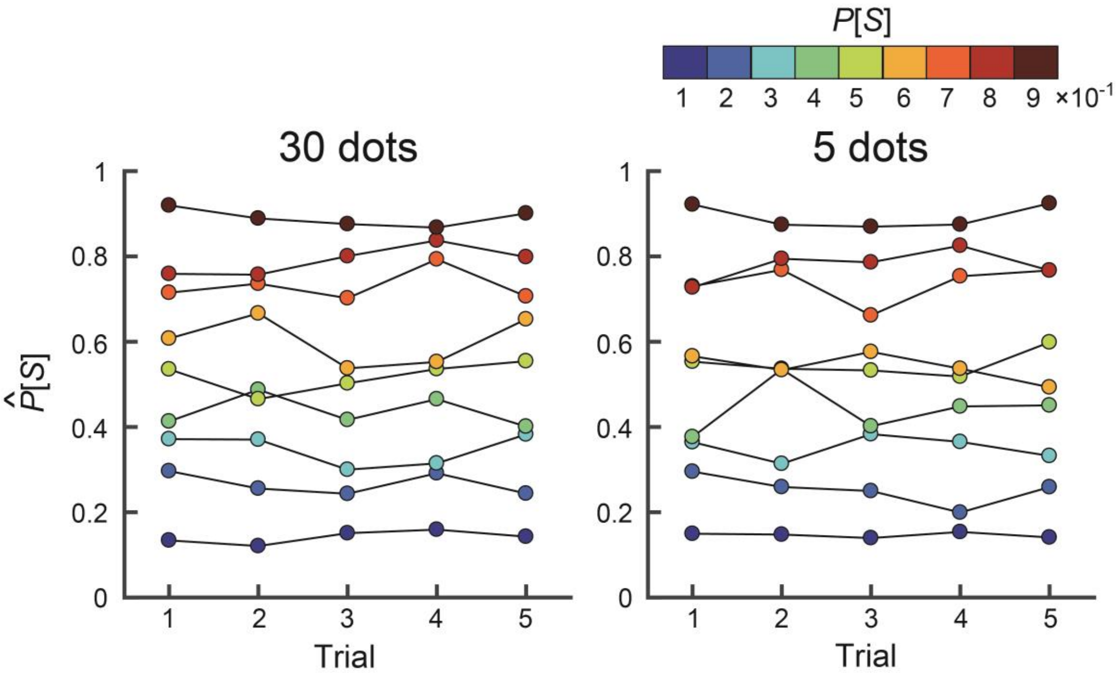

**1-1 Supplementary Figure 1. Estimates of probability did not change systematically across trials.**

The participant's estimates of probability in the symmetric interval are plotted versus trial. Data is averaged across the participants. The color scale of the circle indicates the correct probability between 0.1 and 0.9. The estimates were retained consistently from the beginning to the end of the task. Three-way within-participant ANOVA, using the correct probability (9), sample condition (2), and the number of trials (5) as independent variables, showed no significant main effect of the trial ($F$ [2.6, 40.9] = 1.54, $p$ = 0.22, $\eta^2$ = 0.00).

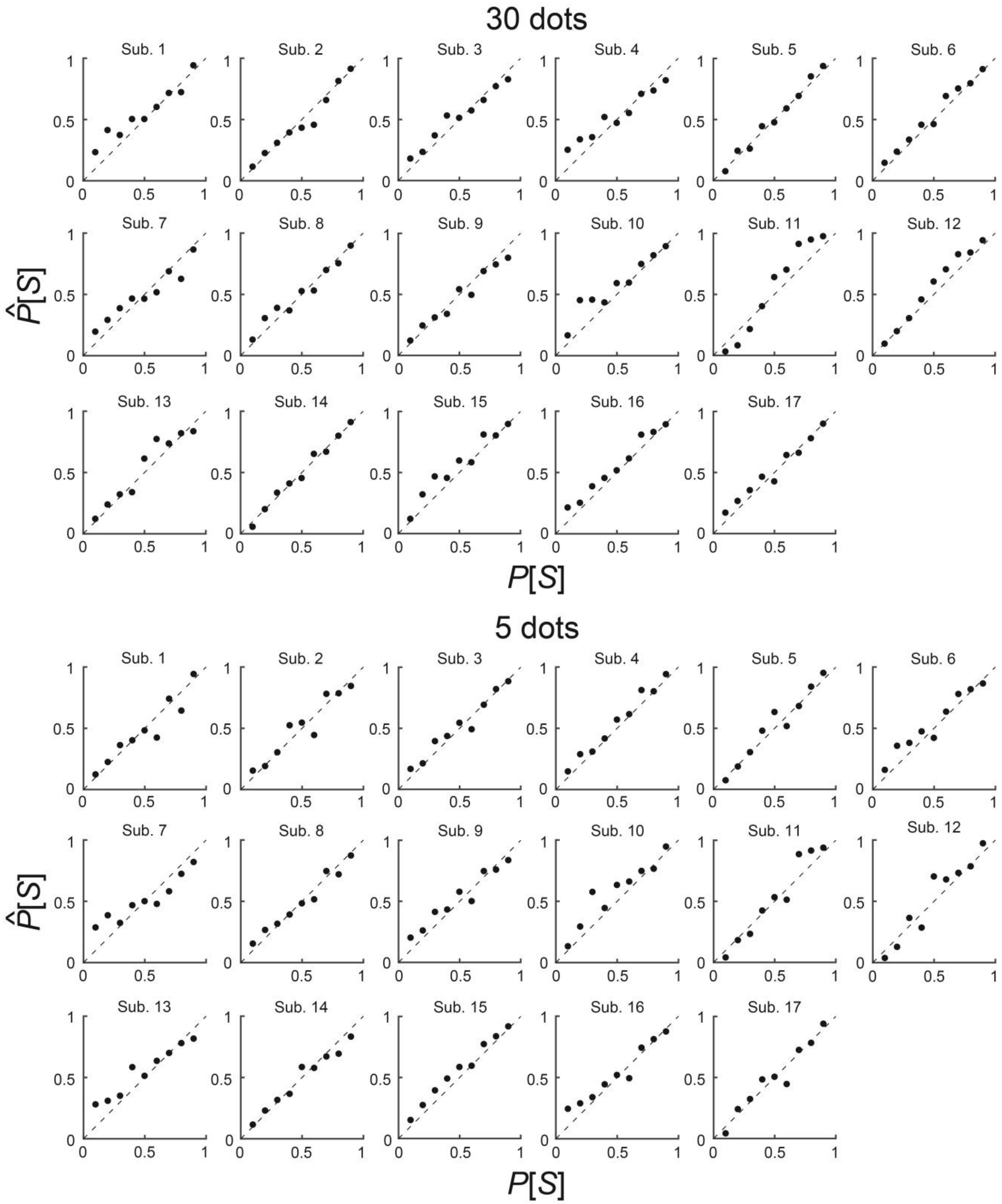

**1-2 Supplementary Figure 2. Plot of accuracy in the symmetric interval for each participant.** The participant's estimates of probability in the symmetric interval are plotted against the correct probability for each participant. Data is averaged across trials.

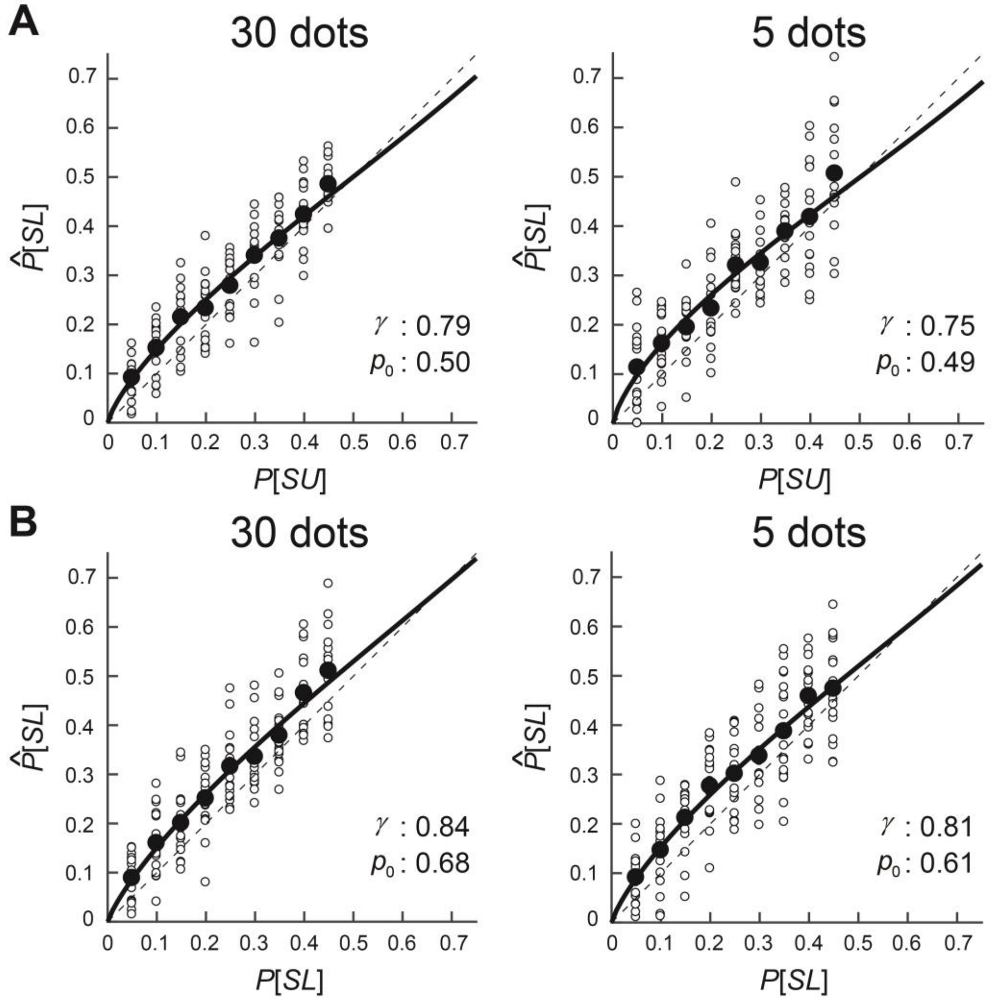

**1-3 Supplementary Figure 3. Plot of accuracy in upper and lower halves of the symmetric interval.**

The participants' estimates of probability in the upper half (**A**) and lower half (**B**) of symmetric interval are plotted against the correct probability. Each white circle denotes the estimates for a single participant and a filled circle is the average estimates across participants. The black thick curve is the best-fit estimate by a linear in log-odds (LLO) function. The LLO parameters for each fit are $\gamma$ and $p_0$.

1-4 Model comparison in the test of accuracy.

We fit participants' estimates in the interval estimation task to the two-parameter linear log-odds function (see Zhang & Maloney, 2012) written as

$$\text{H1:} Lo(\widehat{P}[S]) = \gamma Lo(P[S]) + (1\text{-}\gamma)\, Lo(p_0) \quad \text{[S1]}$$

where $Lo(p) = \ln[p/(1-p)]$. The equation is written in terms of the symmetric interval *S* for convenience. It applies equally to *SU*, *SL* as well. In addition, we considered other hypotheses concerning the form of probability distortion

$$\text{H2:} \widehat{P}[S] = \frac{P[S]^\gamma}{(P[S]^\gamma + (1 - P[S])^\gamma)^{1/\gamma}} \quad \text{[S2]}$$

$$\text{H3:} \widehat{P}[S] = \exp[-(-\text{Ln}(P[S]))^\gamma] \quad \text{[S3]}$$

where hypotheses H2 is a two-parameter version of the probability distortion function from Tversky and Kahneman (1992). H3 is the two-parameter version of the probability distortion function of Prelec (1998). We also compared the LLO function (H1) with a null hypothesis with perfect accuracy (no distortion).

$$\text{H0:} \widehat{P}[S] = P[S] \quad \text{[S4]}$$

We fit these models to the mean estimates across participants by maximum likelihood. Supplementary Table 1 summarizes the results. An AIC model comparison indicates that the LLO function fits the data best.

**1-5 Supplementary Table 1. Model comparison in the test of accuracy.**

| Condition | Model | No. Par. | Fit to the mean estimates | | | | Recovered free parameter(s) | | | |
|---|---|---|---|---|---|---|---|---|---|---|
| | | | 30 | 5 | 30 | 5 | 30 | 5 | 30 | 5 |
| | | | Δi AICc | | Evidence ratio | | $\gamma$ | | $p_0$ | |
| $P[S]$ | H0 | 0 | 0.0 | 0.0 | 1.0 | 1.0 | | | | |
| | H1 | 2 | 12.9 | 1.5 | 629.2 | 2.2 | 0.88 | 0.87 | 0.76 | 0.72 |
| | H2 | 1 | 0.1 | 0.0 | 1.07 | 1.0 | 0.92 | 0.90 | | |
| | H3 | 1 | -2.1 | -1.6 | 0.36 | 0.4 | 0.96 | 0.93 | | |
| $P[SU]$ | H0 | 0 | 0.0 | 0.0 | 1.0 | 1.0 | | | | |
| | H1 | 2 | 14.9 | 8.8 | 1736 | 81.7 | 0.79 | 0.75 | 0.50 | 0.49 |
| | H2 | 1 | 12.3 | 7.8 | 463 | 48.8 | 0.80 | 0.79 | | |
| | H3 | 1 | 6.4 | 4.5 | 25 | 9.6 | 0.77 | 0.74 | | |
| $P[SL]$ | H0 | 0 | 0.0 | 0.0 | 1.0 | 1.0 | | | | |
| | H1 | 2 | 16.0 | 21.0 | 2977.1 | 36527.3 | 0.84 | 0.81 | 0.68 | 0.61 |
| | H2 | 1 | 6.0 | 9.7 | 20.2 | 129.2 | 0.80 | 0.78 | | |
| | H3 | 1 | 2.1 | 4.8 | 2.9 | 11.1 | 0.76 | 0.74 | | |

Note — "No. Par." is the number of free parameters for each model. $\Delta_i \mathrm{AICc} = \mathrm{AICc}_{H1} - \mathrm{AICc}_{Hi}$. A positive AICc difference indicates a better fitting model than H0 (no distortion). The evidence ratio is defined by $\exp(\frac{\Delta_i \mathrm{AICc}}{2})$ and is the relative likelihood of model pairs and represents the evidence about models as to which is better in a K-L information sense (Burnham & Anderson, 1998). A value of evidence ratio means how many times the data is more likely under the alternative than the null hypothesis. Evidence ratios greater than 10 represent strong evidence for the alternative hypothesis whereas evidence ratios less than 0.1 represent strong evidence for the null hypothesis. Evidence ratios greater than 3 (or less than 0.33) represent substantial evidence for the alternative hypothesis (or for the null hypothesis, Jeffreys, 1939/1961). A value in the intermediate range (0.33 and 3) supports neither hypothesis. H1 (LLO function) is strongly supported. The right four columns summarize the free parameters that best describe the data: $\gamma$ is the slope of the curve and $p_0$ is the crossover point.

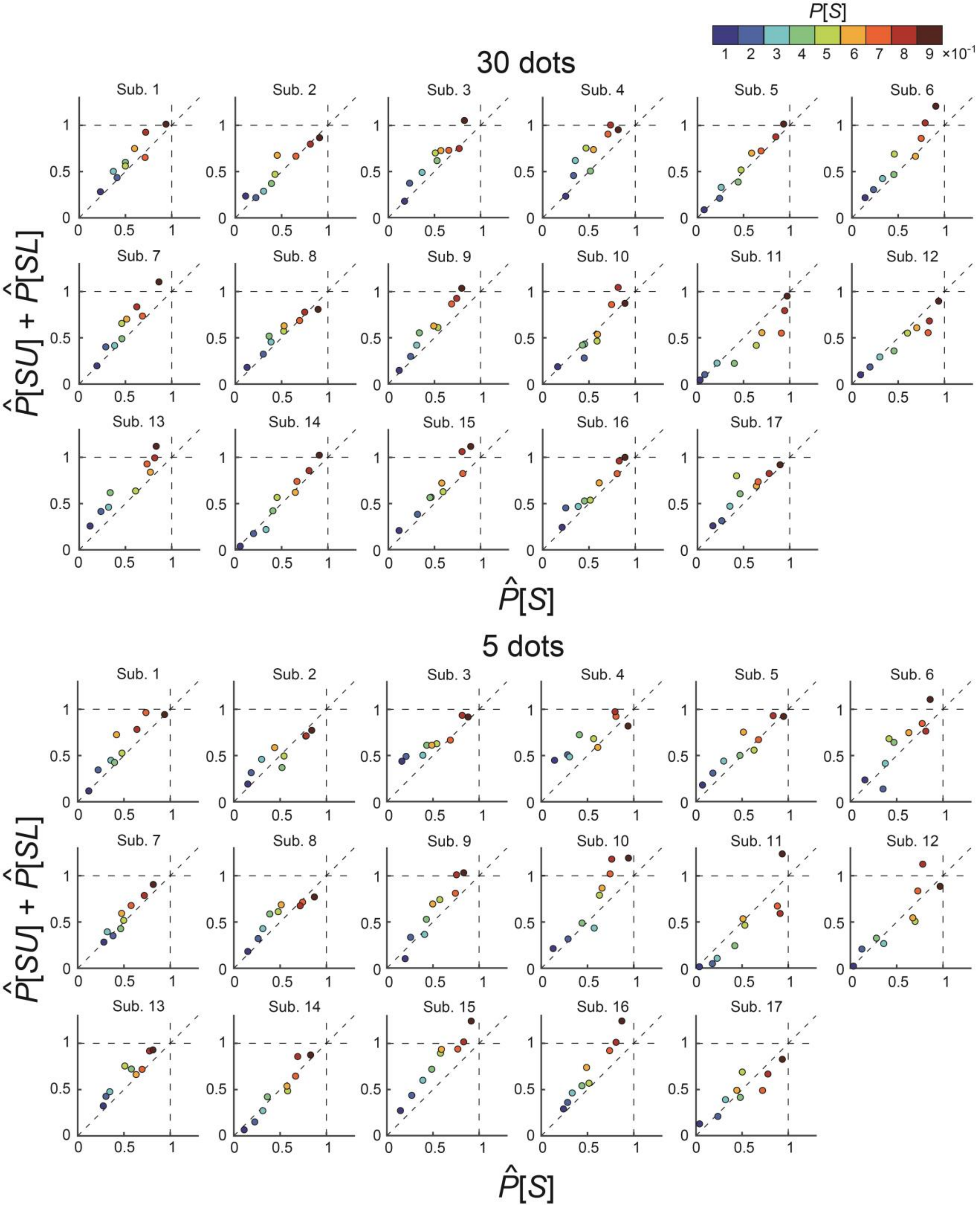

**1-6 Supplementary Figure 4. Plot of additivity in each participant.**

The sum of estimates in upper and lower halves of the symmetric interval are plotted against the estimates of the symmetric interval for each participant. Data is averaged across trials.

**1-7 Supplementary Table 2. Model comparison in the test of additivity.**

| Model | No. Par. | Fit to the mean estimates | | | | Recovered free parameter | |
|---|---|---|---|---|---|---|---|
| | | 30 | 5 | 30 | 5 | 30 | 5 |
| | | Δi AICc | | Evidence ratio | | b | |
| H0 | 0 | 0.0 | 0.0 | 1.0 | 1.0 | | |
| H1 | 1 | 14.8 | 18.3 | 1660 | 9336 | 0.063 | 0.072 |
| H2 | 1 | -2.6 | -2.6 | 0.3 | 0.3 | 0.000 | 0.000 |

See the note in Supplementary Table 1. $\Delta_i \mathrm{AICc} = \mathrm{AICc}_{H0} - \mathrm{AICc}_{Hi}$. A positive AICc difference indicates a better fitting model than a null hypothesis (H0). Three hypotheses concerning estimates for the test of additivity are below. A super-additive model (H1) outperformed the other models.

$$\mathrm{H0}: \widehat{P}[SU] + \widehat{P}[SL] = \widehat{P}[S] \quad \text{[S5]}$$

$$\mathrm{H1}: \widehat{P}[SU] + \widehat{P}[SL] = \widehat{P}[S] + b\,(b > 0) \quad \text{[S6]}$$

$$\mathrm{H2}: \widehat{P}[SU] + \widehat{P}[SL] = \widehat{P}[S] + b\,(b < 0) \quad \text{[S7]}$$

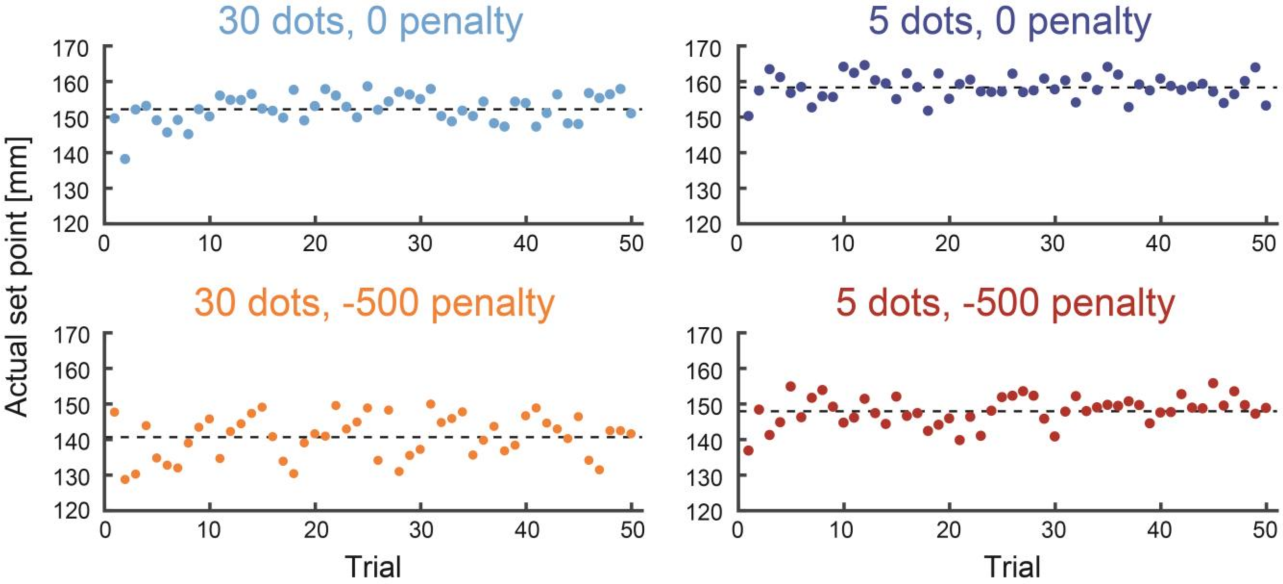

**1-8 Supplementary Figure 5. Trial-by-trial set point in the decision task.**

Trial-by-trial set points averaged over the participants is plotted for each condition. The horizontal dashed lines denote the mean set point across all trials. There is no evident pattern in the residuals.

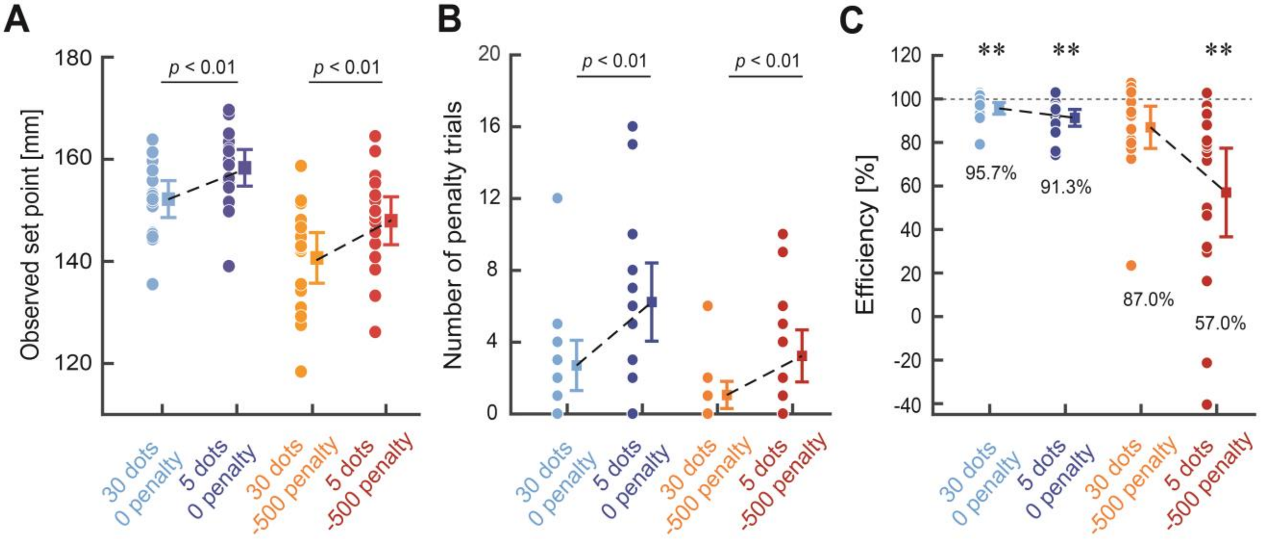

**1-9 Supplementary Figure 6. Performance index in the decision task.**

**A.** The decision makers' actual set points. Each circle denotes the individual data averaged across trials, and a rectangle denotes the average across all participants. The error bars indicate ± 2 s.e.m. **B.** The number of penalty trials that the participants incurred in each condition. The error bars indicate ± 2 s.e.m. A two-way within-participant ANOVA showed a main effect of the penalty condition ($F$ [1, 16] = 91.78, $p$ = 0.001, $\eta^2$ = 0.20) and a main effect of the number of dots ($F$ [1, 16] = 22.73, $p$ = 0.001, $\eta^2$ = 0.30). The participants thus incurred a larger number of penalties with fewer samples and with a smaller penalty. **C.** The efficiency, as the ratio of the actual total score to the maximum total score possible predicted by the normative decision model. The actual total scores were significantly smaller than the maximum possible scores in all conditions except the 30 dot, -500 penalty condition (two-tailed paired-sample $t$-test: $ts$ [16] > 3.25, $ps$ < 0.005, $ds$ > 1.15, Bonferroni corrected for four conditions). ** indicates $p$ < 0.01 from the maximum possible score.

#### 2 Supplementary methods

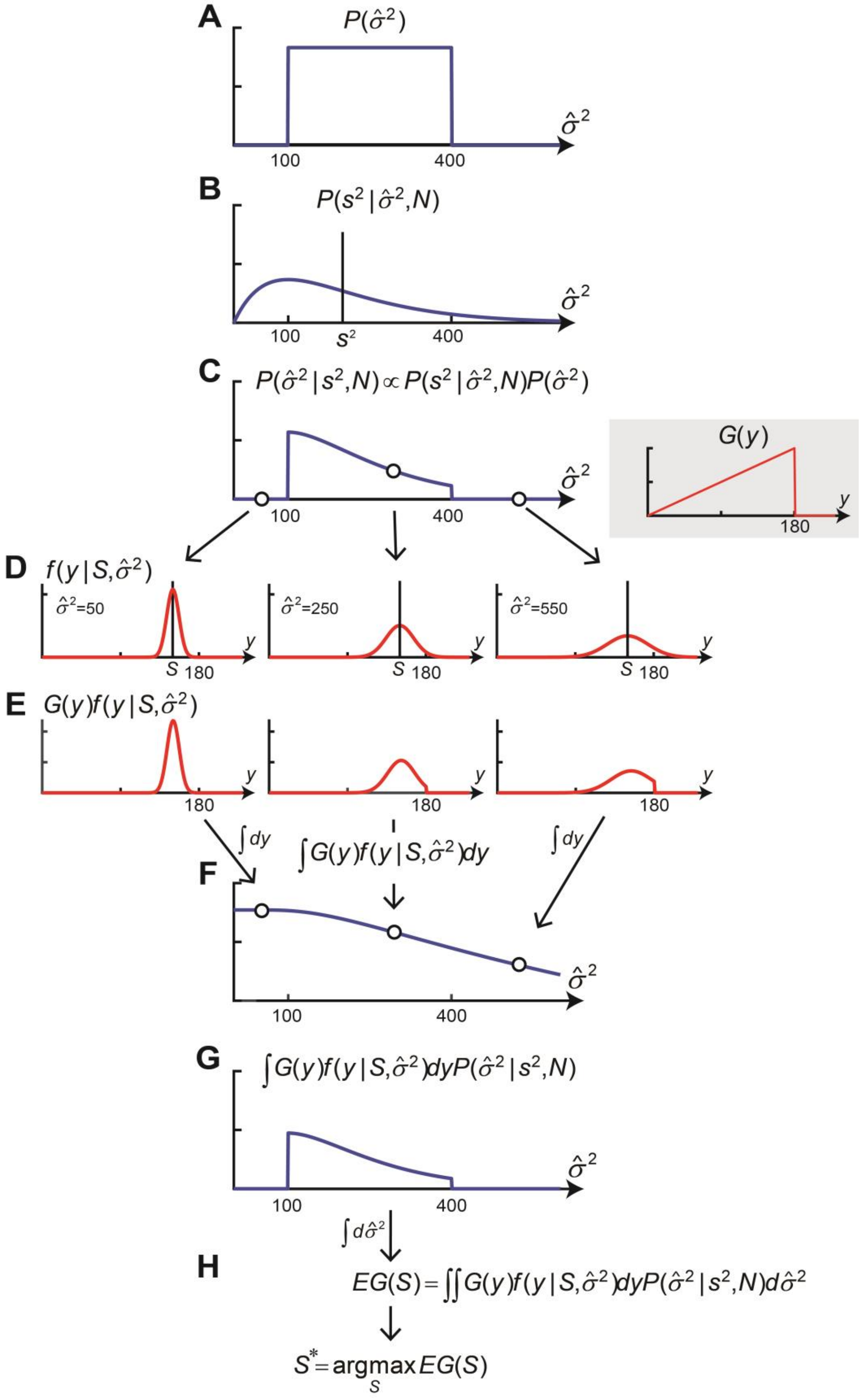

**2-1 Supplementary Figure 7. Illustration for the procedure of the sufficient model.**

**A.** Prior distribution of the estimates of population variance $\hat{\sigma}^2$. On each trial, the population variance was uniformly chosen from the range between $\sigma^2 = 100\text{mm}$ and $\sigma^2 = 400\text{mm}$. The prior distribution was set to be the same distribution for generating the population variance. **B.** Likelihood function of the estimates of population variance. Given the number of points in the sample $N$ and the sample variance $s^2$, the likelihood function of the estimates of population variance as a chi-square distribution with $N - 1$ degree of freedom can be estimated. A solid vertical line shows the sample variance on this trial. We set the number of points to $N = 5$. **C.** Posterior probability distribution of the estimates of population variance. The posterior is a product of the prior and likelihood function. **D.** Estimated probability density function modeled as a Gaussian distribution. The width of the pdf depends on the estimated population variance. We show three example estimated pdfs. In the decision task, the location of the estimated pdf can be shifted by a set point $S$ (here we chose $S = 150\text{mm}$). A small gray inset shows a reward function $G(y)$ . **E.** The function illustrates the product of the estimated pdf with the reward function. **F.** The ideal decision maker integrates the estimated pdf with the reward function given the set point and the estimated population variance, which produces the expected reward function as a function of the estimated population variance. **G.** The ideal decision maker then scales the expected reward function **F** by the posterior probability of the estimated population variance **C**. **H.** The final output of the expected reward can be obtained by integrating the function shown in **G** over $\hat{\sigma}^2$. The ideal decision maker chooses the ideal set point $S^*$ maximizing expected reward can be maximized.

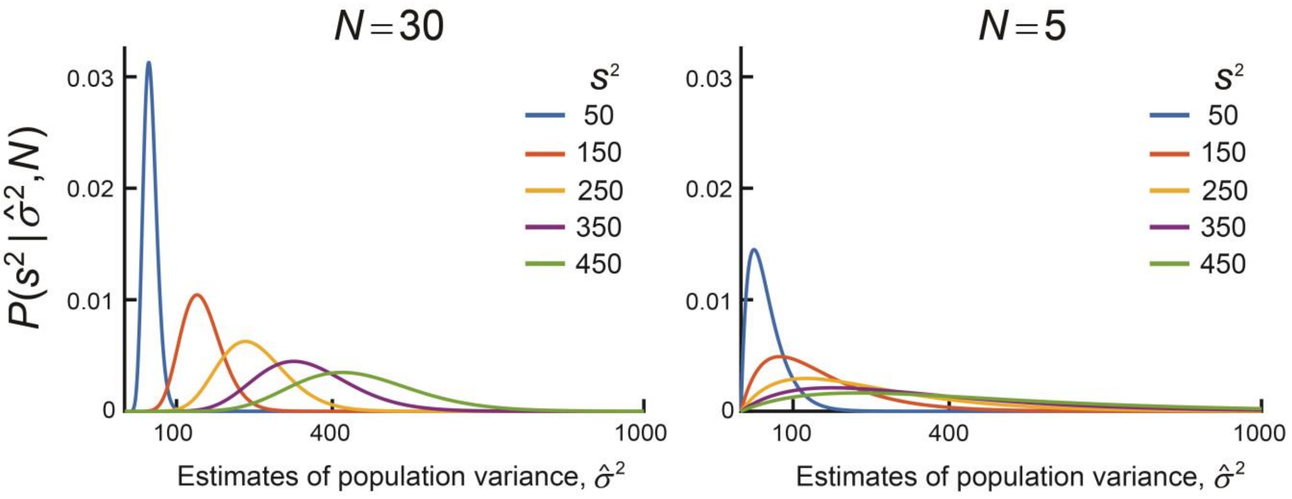

**2-2 Supplementary Figure 8. Likelihood function of the estimated population variance as a chi-square distribution.**

Given the population variance $\hat{\sigma}^2$, the random variable of the sample variance $s^2$ is distributed according to a chi-square distribution with $N-1$ degrees of freedom. Therefore, the likelihood function of the estimated population variance can be described as a chi-square probability density function. We show the likelihood functions when $N=30$ (left) and $N=5$ (right). The sample variance varied between 50 mm and 450 mm.
